# Rapid dynamics of electrophysiological connectome states are heritable

**DOI:** 10.1101/2024.01.15.575731

**Authors:** Suhnyoung Jun, Stephen M. Malone, William G. Iacono, Jeremy Harper, Sylia Wilson, Sepideh Sadaghiani

## Abstract

Time-varying changes in whole-brain connectivity patterns, or connectome state dynamics, are a prominent feature of brain activity with broad functional implications. While infra-slow (<0.1Hz) connectome dynamics have been extensively studied with fMRI, rapid dynamics highly relevant for cognition are poorly understood. Here, we asked whether rapid electrophysiological connectome dynamics constitute subject-specific brain traits and to what extent they are under genetic influence. Using source-localized EEG connectomes during resting-state (N=928, 473 females), we quantified heritability of multivariate (multi-state) features describing temporal or spatial characteristics of connectome dynamics. States switched rapidly every ∼60-500ms. Temporal features were heritable, particularly, Fractional Occupancy (in theta, alpha, beta, and gamma bands) and Transition Probability (in theta, alpha, and gamma bands), representing the duration spent in each state and the frequency of state switches, respectively. Genetic effects explained a substantial proportion of phenotypic variance of these features: Fractional Occupancy in beta (44.3%) and gamma (39.8%) bands and Transition Probability in theta (38.4%), alpha (63.3%), beta (22.6%), and gamma (40%) bands. However, we found no evidence for heritability of spatial features, specifically states’ Modularity and connectivity pattern. We conclude that genetic effects strongly shape individuals’ connectome dynamics at rapid timescales, specifically states’ overall occurrence and sequencing.

## 1. Introduction

Investigations of the functional connectome have been dominated by fMRI due to its superb spatial resolution, which however does not allow the study of rapid connectome dynamics at sub-second, cognitively highly relevant timescales. A growing body of work has established that non-invasive, real-time methods, i.e., EEG and MEG, can provide a window into the connectome and, importantly, its rapid dynamics (static/time-averaged connectome: (de Pasquale et al., 2010; Brookes et al., 2011; Deligianni et al., 2014; Hipp and Siegel, 2015; Wirsich et al., 2017), connectome dynamics: (Baker et al., 2014; Brookes et al., 2014; Sitnikova et al., 2020; Wirsich et al., 2020; Coquelet et al., 2022). EEG/MEG connectome studies capitalize on methodological advances that combine source-localization with correction of source leakage, an artifact arising from the activity of the same electrical source in the brain being picked up by multiple sensors. As such, real-time methods can extend our understanding of connectome dynamics beyond the knowledge gained from fMRI (for review, see (Sadaghiani and Wirsich, 2020)).

What we know from fMRI connectome studies suggests that the time-varying dynamics in large-scale connectivity are of functional importance for cognition (Preti et al., 2017; Cohen, 2018). These reconfigurations can be characterized as flexible changes in connectome states, representing varying strengths of connectivity between specific sets of brain regions within the whole-brain connectome, which occur repeatedly over time. Such time-varying features have been linked to a wide range of behaviors and cognitive processes, encompassing both the temporal organization of connectome state transitions (Vidaurre et al., 2017; Eichenbaum et al., 2020; Jun et al., 2022) and changes in the spatial organization of connectome states (Thompson et al., 2013; Sadaghiani et al., 2015; Douw et al., 2016). Particular temporal features of fMRI-derived connectome dynamics, specifically the proportion of the total recording time spent in each connectome state (Fractional Occupancy) and the probability to transition between specific pairs of connectome states (Transition Probability), have been linked to behavioral performance (Vidaurre et al., 2017; Eichenbaum et al., 2020; Jun et al., 2022) and found to be heritable (Vidaurre et al., 2017; Jun et al., 2022). More specifically, our previous fMRI work has established substantial genetic effects (*h^2^* ∼ 40%) and behavioral relevance of Fractional Occupancy and Transition Probability (Jun et al., 2022). We further identified specific genetic polymorphisms predictive of fMRI-derived Fractional Occupancy and Transition Probability via the regulatory impact of modulatory neurotransmitter systems (Jun et al., *Under Review;* https://doi.org/10.17605/OSF.IO/VF2ZW).

However, as mentioned above, the temporal dynamics captured by the slow fMRI-derived indirect measure of neural activity, or BOLD signal, limit the study of rich, sub-second temporal dynamics. While direct electrophysiological techniques, i.e., MEG or EEG, can capture such rapid dynamics, their use in the study of individual differences and heritability has primarily focused on the electrophysiological power spectrum or *static* (time-averaged) connectivity (Posthuma et al., 2005; Smit et al., 2005; Colclough et al., 2017). When investigating large-scale brain dynamics with M/EEG, one area of focus has been microstates, denoting recurrent, spatially diffuse sensor-level topographies that transition rapidly (every ∼40-200 ms) (Lehmann et al., 2009; Coquelet et al., 2022). These EEG microstates were found to be predictive of interindividual variability in cognitive abilities (Muthukrishnan et al., 2016; Kim et al., 2021) and linked to neurodegenerative and psychiatric disorders (Hatz et al., 2015; da Cruz et al., 2020). However, the topographies reflected in microstates are typically spatially diffuse, because they are estimated in “sensor space” rather than on the cortical surface and thus fall short of informing about connectome states that encompass spatially localized coactivations across networks of brain regions. Recently, we have shown that rapid temporal dynamics of recurrent connectome states are of functional significance, explaining individuals’ cognitive abilities (Jun et al., 2024; Submitted). This individual specificity invites the question to what degree such rapid connectome state dynamics are heritable.

In the current study, we investigated whether temporal and spatial features of fast connectome dynamics in canonical electrophysiological frequency bands are attributable to genetic effects. To address this question, we applied Hidden Markov Modelling (HMM) to extract discrete brain states using source-reconstructed resting-state EEG data from the Minnesota Twin Family Study (Keyes et al., 2009; Wilson et al., 2019). The dataset included monozygotic (MZ) and dizygotic (DZ) twin pairs, and pairs of unrelated individuals. Consistent with our preceding heritability study in fMRI (Jun et al., 2022), we included two temporal features and two spatial features based on prior evidence for their behavioral relevance. Interestingly, only temporal features were found to be heritable in the fMRI study. Yet, we included the spatial features in the current study given that rapid EEG-derived dynamics might capture connectivity processes different from those observed in fMRI. The temporal features consisted of Fractional Occupancy and Transition Probability, and the spatial features included time-varying Modularity (Modularity_Time-Varying_) and the time-varying strength of functional connectivity (FC_Time-Varying_) of the set of connections (clusters) exhibiting the strongest cross-state change. We examined the effect of genetic relatedness on each multi-variate connectome feature and fitted quantitative genetic models to quantify the genetic effects. To the best of our knowledge, this is the first EEG study investigating heritability of source-space connectome dynamics. This study sheds light on genetic contributions to individual differences in sub-second connectome dynamics.

## 2. Materials and Methods

Figure 1 is a schematic representation of the overall approach and analysis subsections.

**Figure 1.**
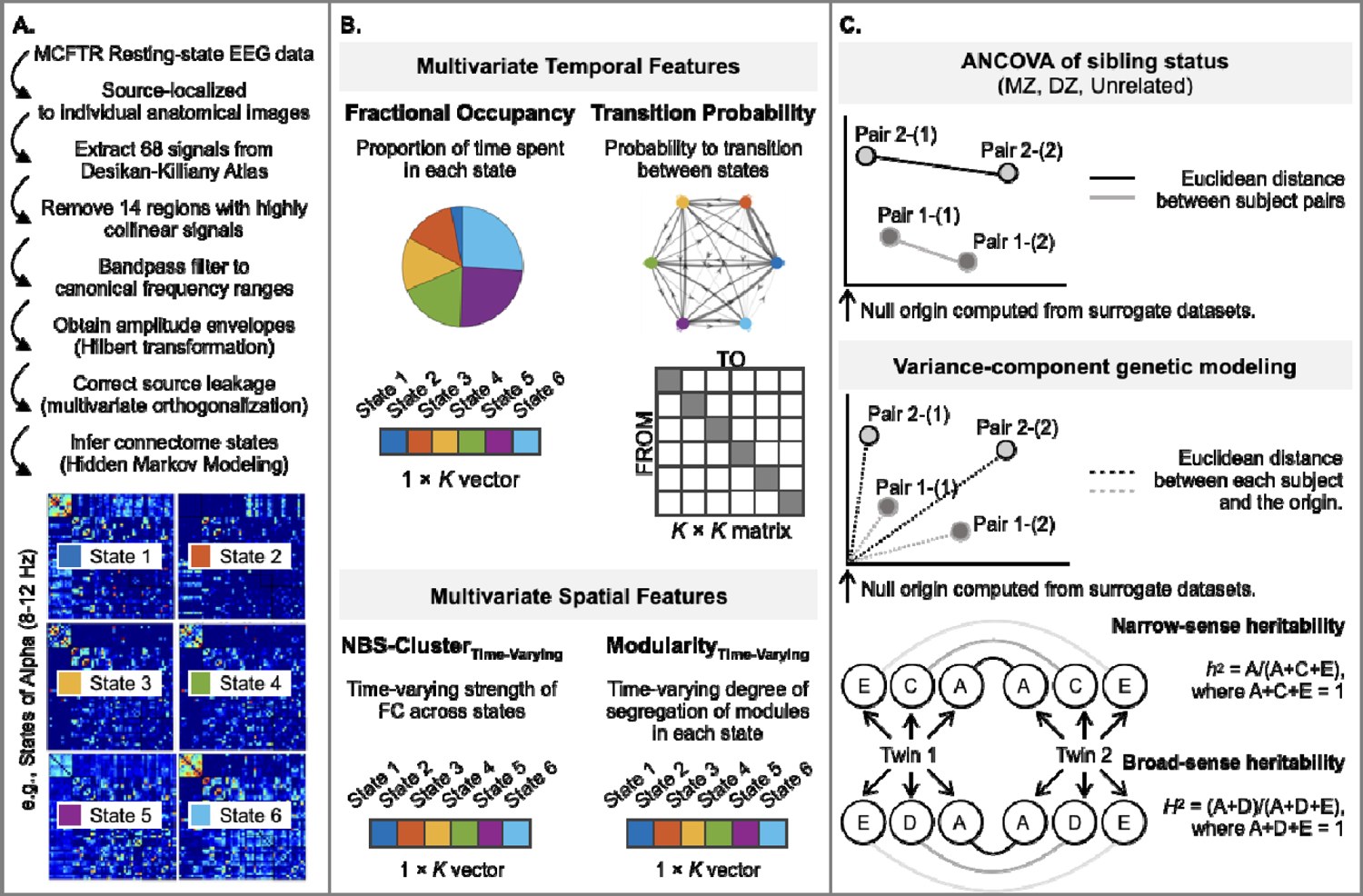
An overview of the analysis pipeline. [A]. We used resting-state source-space EEG and structural images from Minnesota Twin Family Study samples. We employed a hidden Markov model (HMM) to extract *K*=6 discrete connectome states (or *K*=4 states for replication) for each canonical EEG frequency band associated with state time course for each subject indicating the probability of when each state is active. The amplitude coupling-based FC matrices of the states are shown at the bottom. The six states are color coded (blue, red, yellow, green, purple, and light blue) to illustrate their contribution to the connectome’s dynamic features of interest. **[B]** We constructed each feature in a multivariate manner to comprehensively represent all states. Specifically, multivariate temporal features were defined as the proportion of the recording time spent in each connectome state (Fractional Occupancy) and the probability matrix of transitioning between all possible pairs of discrete states (Transitio Probability). Multivariate spatial features include time-varying Modularity (Modularity_Time-Varying_), and time-varying connectivity strength (FC_Time-Varying_) averaged across the set of connections (region-pairs) with the strongest dynamic changes across states (Zalesky et al., 2010). **[C]** We tested whether genetically more related subjects displayed greater similarity in their multivariate features than genetically less related subjects. First, for each feature of dimension *m*, we estimated a null model-derived origin point in the *m*-dimensional space. The position of eac subject’s multi-dimensional feature value was estimated relative to this origin for genetic modeling. Further, the similarity of this position between a given pair of subjects was quantified as Euclidean distance for ANCOVA analyses; a one-way ANCOVA of the factor sibling status with three levels (monozygotic twins (MZ), sex-matched dizygotic twins (DZ), and sex-matched pairs of unrelated individuals) was performed on the distance value for each of the features. Secondly, we employed structural equation modeling (i.e., genetic variance component model) to quantify the genetic effects. Phenotypic variance of a trait was partitioned into additive genetic (denoted A), common environmental (denoted C) Or dominant genetic effect (denoted D) and unique environmental components (denoted E). Narrow-sense heritability (*h^2^*) is quantified as the proportion of variance attributed to the genetic factor (A) and broad-sense heritability (*H^2^*) as the proportion of variance attributed to A and D factors.

### 2.1. Subjects

Participants for the present investigation are from the two independent cohorts of twins from the Minnesota Twin and Family Research (MCTFR) (Iacono et al., 1999; Keyes et al., 2009; Wilson et al., 2019). Twins in both cohorts have been followed periodically since approximately the age of 11. As part of their most recent assessment, participants underwent structural MRI scans in addition to resting EEG recordings. At time of initial recruitment and at each follow up, participants gave written informed consent or assent, if under the age of 18 for their participation.

A total of 1164 subjects in two independent cohorts of MCTFR twins completing virtually identical assessments, 928 had usable and complete EEG data which was source-localized successfully (see **2.3 EEG signal pre-processing and source localization**), thus permitting HMM-based estimation of discrete connectome states. The included subjects (473 females) were 23–40 years of age at time of data acquisition. Subsequently, 463 sex-matched pairs (926 subjects) were formed for heritability analysis as follows: 206 monozygotic (MZ) twin pairs, 112 sex-matched dizygotic (DZ) twin pairs, and 145 pairs of sex-matched unrelated individuals. Each subject entered only one pair. Note that the unrelated individuals are twins whose co-twin lacked complete data and was therefore not part of the analytic sample.

### 2.2 MRI and EEG acquisition

Structural MRI data were collected on either 3T Siemens Trio or Prisma MRI scanner (32-channel array head coil) at the Center for Magnetic Resonance Research, University of Minnesota. Three-dimensional T1-weighted sagittal plane anatomical images were acquired using the following magnetization-prepared rapid gradient echo sequence: TR = 2530 ms; TE = 3.65 ms; flip angle = 7°; matrix size = 256 × 256; FOV = 256 mm; GRAPPA = 2; 240 coronal slices with 1-mm isotropic voxels; single shot; interleaved acquisition.

While recording EEG, participants rested comfortably in a darkened room, with their head and neck supported while hearing 55-dB white noise played through headphones. They were instructed to keep their eyes closed and relax. A recorded voice subsequently instructed them to open the eyes or close them at 1-min intervals. A total of 6 min of EEG was collected, 3 min with eyes open and 3 with eyes closed. EEG data were acquired from 61 scalp electrodes arranged according to the International 10/10 system using a BioSemi ActiveTwo system (BioSemi, Amsterdam, The Netherlands) at 1024 Hz. ActiveTwo amplifiers are DC coupled. ActiveTwo signals are monopolar. They were low-pass filtered using a digital 5th-order Bessel antialiasing sinc filter with a cutoff frequency (3-dB attenuation) of 205 Hz. Pairs of electrodes placed above and below the right eye or on the outer canthus of each eye allowed for detecting blinks and other eye movements. Additional electrodes were placed on left and right earlobes, and the average of these signals was derived offline to serve as a reference.

### 2.3. EEG signal pre-processing and source localization

Raw resting-state EEG signals were pre-processed using a monitored automated pipeline (https://www.github.com/sjburwell/eeg_commander) of the Minnesota Twin Family Study group and EEGLAB (Delorme and Makeig, 2004) in MATLAB (version R2021b, Mathworks, Inc.). Signals were down-sampled to 256 Hz, filtered with a 0.1 Hz high-pass filter (*firfilt* EEGLAB plugin; 1,286 Kaiser window), and referenced to averaged earlobe signals. A monitored automated pipeline detected four kinds of signal anomalies: disconnected channels/flat signals, interelectrode electrolyte bridging (Tenke and Kayser, 2001), large amplitude deviations, and muscle/cap shift (motion) noise. Descriptives (e.g., temporal variance) were calculated for each electrode and 1-s time range. Data that exceeded four normalized median absolute deviations from the median (Rousseeuw and Croux, 1993) in 25% of a 1s time range or 75% of a given electrode were removed. Among others, this approach is effective in removing periods with head motion artifacts. Ocular correction was performed using independent components (IC) analysis (infomax algorithm; Bell and Sejnowski, 1995) and joint consideration of temporal and spatial signal characteristics. The IC time series and inverse weights were compared with the time courses of the bipolar vertical or horizontal EOG and the inverse weight of a stereotypical blink or horizontal saccade to correct for vertical and horizontal ocular artifacts, respectively. If the squared joint temporal and spatial correlations for an IC exceeded an empirically calculated threshold based on Expectation Maximization (Mognon et al., 2011), that IC was subtracted from the data.

For source localization, we imported preprocessed EEG recordings and MR-based anatomical images into Brainstorm software (Tadel et al., 2011). The EEG signals were resampled to 250 Hz, corrected for DC offsets, linearly detrended, and low-pass filtered at 70 Hz. We manually marked fiducial points, including the anterior commissure (AC), posterior commissure (PC), inter-hemispheric point, nasion (NAS), and left and right pre-auricular points (LPA and RPA), of all subjects using their individual anatomical images to aid coregistration of electrode positions and T1 images. The coregistration was refined by manually moving the electrode positions onto the electrode artifacts visible in the T1 image. We then used the OpenMEEG software (Gramfort et al., 2010) with a symmetric boundary element method (BEM) to calculate a forward model of the skull based on the individual T1 image of each subject (Tadel et al., 2019). Then, we used the Tikhonov-regularized minimum-norm estimation (MNE) inverse method to compute the sources, with default parameter settings for regularization and source depth weighting (Tikhonov parameter = 10%, assumed SNR = 3.0, constrained sources normal to cortex, depth weighting 0.5/max amount 10) (Baillet et al., 2001; Tadel et al., 2019).

### 2.4. Parcellation and Source-leakage correction

We used the Desikan-Killiany Atlas (Desikan et al., 2006) in Brainstrom to average source signals within each of the atlas’ 68 anatomically distinct brain regions. To aid network-level interpretation, we also determined each region’s membership within the seven canonical Intrinsic Connectivity Networks or ICNs (Yeo et al., 2011) based on spatial overlap.

To mitigate source-leakage confounds caused by the blurring of point dipole sources and the spreading of signals across neighboring regions, we excluded regions whose signals were collinear with others based on a QR decomposition, which is commonly used to model the correlation structure of a set of variables, using the *qr* function in Matlab. As a result, 14 regions were excluded from the investigation (*Figure S1*). The remaining 54 regional signals underwent detrending and bandpass filtering within canonical frequency ranges: delta (1-3 Hz), theta (4-7 Hz), alpha (8-12 Hz), beta (13-25 Hz), and gamma (30-45 Hz). Then, we used a symmetric orthogonalization procedure (Colclough et al., 2015) to remove all shared signal at zero lag between the regions. This multivariate method extends previous orthogonalization methods (Brookes et al., 2012; Hipp et al., 2012) and identifies orthogonal time-courses that maintain the closest similarity to the original, unmodified time-series. Finally, amplitude envelopes for each canonical frequency band and brain region were computed using the Hilbert transform, which were then downsampled to 40 Hz (Baker et al., 2014; Hunyadi et al., 2019).

### 2.5. Hidden Markov Modelling of connectome states

The hidden Markov model (HMM) assumes that time series data can be represented by a finite sequence of hidden states. Each HMM-inferred connectome state, along with its corresponding time series, represents a unique connectivity pattern that re-occurs over time. Using the HMM-MAR toolbox (Vidaurre et al., 2016), we applied the HMM to the region-wise EEG amplitude timeseries separately for each frequency band to derive discrete recurrent connectome states characterized by their mean activation and FC matrix. We obtained six connectome states (*K* = 6). While HMMs require an *a priori* selection of the number of states, *K*, the objective is not to establish a ‘correct’ number of states but to strike a balance between model complexity and model fit and to identify a number that describes the dataset at a useful granularity (Quinn et al., 2018). Our previous fMRI-based investigation into connectome heritability (Jun et al., 2022) reported results for two different *K* values (to ensure that outcomes are not limited to a single chosen parameter), namely *K* of 4 and *K* of 6. This choice was in turn informed by prior fMRI literature (Vidaurre et al., 2016; Karapanagiotidis et al., 2020). The choice of *K =* 4 and 6 falls with the range applied in prior HMM studies of EEG and MEG data, which have used *K*s between 3 and 16 (Baker et al., 2014; Vidaurre et al., 2016; Quinn et al., 2018; Hunyadi et al., 2019; Coquelet et al., 2022), where two of the studies used *K* of 6. Therefore, based on the success of our prior fMRI study in revealing heritability within 6-state and 4-state models (Jun et al., 2022), the current study reports results from *K* of 6 (main text) and *K* of 4 (supplementary materials).

### 2.6. Null model of Hidden Markov Models

To demonstrate that the dynamic trajectory of connectome state transitions is not occurring by chance, we employed a null model. This involved generating 50 simulated state time courses for each frequency band, which were of the same length as the original empirical state time courses. While preserving the static covariance structure, the temporal ordering of states was intentionally disrupted (Vidaurre et al., 2016). It is worth noting that selecting 50 simulations for each of the frequency bands in this analysis represents a rigorous choice in comparison to previous studies (e.g., four simulations in (Vidaurre et al., 2017)). We performed HMM inference with *K*s of 6 (and 4 for replication) on each of these simulated time courses, allowing us to recalculate all above-described temporal and spatial connectome features at both the group and subject levels. Through this process, we confirmed that the original dataset’s non-random distribution of features over states represented veridical dynamics as it was absent in the simulated data (*Figure S2*). We further used the surrogate data for heritability testing as detailed below.

### 2.7. Multivariate temporal features of the dynamic connectome

The HMM-derived estimates provide a comprehensive set of multivariate temporal features that simultaneously characterized all states of the dynamic connectome. These estimates describe the temporal aspects of connectome dynamics by characterizing the sequence of connectome states, namely the trajectory of the connectome through state space. For each subject, we calculated the Fractional Occupancy (the proportion of total time spent in a given state; 1 × *K*) and Transition Probability (the probability matrix of transitioning between all possible pairs of discrete states; *K* × *K*). Notably, our previous work demonstrated strong genetic effects specifically on these two temporal features in fMRI-derived functional connectomes (Jun et al., 2022).

### 2.8. Multivariate spatial features of the dynamic connectome

In line with our prior fMRI study (Jun et al., 2022), we also incorporated several multivariate spatial features to describe the functional connectivity (FC) arrangement of states. While no spatial features were found to be heritable in the fMRI study, outcomes might differ for rapid EEG-derived dynamics. To assess the level of segregation for each connectome state, we estimated Newman’s Modularity (Newman, 2006), a fundamental global topological characteristic. The Brain Connectivity Toolbox (Rubinov and Sporns, 2010) was employed to quantify Modularity, where the modular partition was configured to comprise the canonical ICNs (Yeo et al., 2011). The Modularity value for the *K* states were then combined into a *K*-dimensional vector constituting the multivariate feature (Modularity_Time-Varying_) for heritability analysis.

Our second spatial feature was derived from clusters of connections that exhibited significant differences in FC strength across the *K* connectome states. As an initial step, we conducted mass univariate *F*-tests across states for all connections, adjusting for age and sex. For each connection, the resulting (absolute) *F*-value reflects its change in connectivity value across the *K* states. Subsequently, upon thresholding *F*-values at a variety of connection densities (1-5%), we performed the Network-Based Statistics (NBS) permutation method to identify sets of connected edges or clusters that showed significant differences at family-wise error rate corrected *P* < .05 (see *Figure S6* for visualization of binary matrix of a data-driven set of clusters of connections). At each density, the edge-wise FC values within the identified clusters were averaged separately for each of the *K* states, resulting in a 1 x *K* vector representing the time-varying FC (FC_Time-Varying_) of the data-driven clusters for heritability analysis. In the main manuscript, we present results for a 5% density and provide generalizations to other densities in *Tables S3*.

For exploratory analyses, we expanded our scope by incorporating a more comprehensive collection of multivariate spatial features. Specifically, we examined FC_Time-Varying_ between all pairs of the seven canonical ICNs, including the within-network connectivity of each ICN. For each ICN pair and each of the *K* states, we averaged FC values among their connections and combined them into a *K*-dimensional vector. This vector represents the multivariate feature that encompasses the FC_Time-Varying_ of the corresponding ICN pair for the purpose of heritability analysis (see *Table S4* for details).

### 2.9. Similarity estimation and heritability testing

The procedures carried out in our previous twin heritability study (Jun et al., 2022) laid the groundwork for the current investigation. Initially, for each multivariate connectome dynamics feature and separately for each frequency band, we constructed a multidimensional space by setting the origin point as the average of the feature from the 50 surrogate datasets, as described in the **null model (2.6)** section above. Using the multidimensional space, we calculated the pairwise similarity of each feature by measuring the Euclidean distance between pairs of subjects (Colclough et al., 2017; Jun et al., 2022). Crucially, this similarity estimation approach preserved the positional relationship between elements in each multivariate feature. To investigate the relationship between the genetic makeup of subject pairs and the similarity of each multivariate feature, we conducted a one-way analysis of covariance (ANCOVA) of sibling status (MZ twins, sex-matched DZ twins, and sex-matched unrelated individuals) on the similarity of each multivariate feature, adjusting for the difference in age and sex between the subject pairs.

Additionally, we explored whether the heritability of attributes was driven by the overall pattern or by specific components (i.e., state-by-state elements) of each multivariate feature. Specifically, we assessed the similarity of each state-specific component of the Fractional Occupancy, FC_Time-Varying_ of data-driven clusters and Modularity_Time-Varying_ between a given pair of subjects. Likewise, we estimated the similarity of each off-diagonal state-pair component of the Transition Probability matrix. To examine the effects of sibling status and connectome state on the similarity of individual components of the multivariate features, we employed two-way ANCOVAs of the factors sibling status and connectome state.

Finally, we employed structural equation modeling, commonly used in classical twin studies, to quantify the variance in dynamic connectome features explained by genetic effects. This modeling approach, with underlying biological assumptions (Keller and Coventry, 2005), partitions the phenotypic variance into the three distinct components using maximum likelihood methods: additive genetic variance (A), accompanied by either common environmental variance (C) or dominant genetic variance (D), and random environmental variance (E) (Yashin and Iachine, 1995). Leveraging these components, heritability is computed as the portion of phenotype variance explained by genetic variance, denoted as narrow-sense heritability (*h^2^*) in the ACE model and broad-sense heritability (*H^2^*) in the ADE model.

The genetic variance model requires each subject to have a singular value for each phenotype to estimate the correlation of univariate phenotypes among twin pairs. To accommodate our multivariate phenotypes within this framework, we computed the Euclidean distance for each subject’s multivariate phenotype from the origin point which we established (Figure 1). To implement this method, we utilized the R package *mets* (http://cran.r-project.org/web/packages/mets/index.html), adjusted for age and sex. Subsequently, we employed nested models, namely AE, CE, or DE, to gauge the statistical significance of these nested structures using a likelihood ratio test and assessed the fitness of each model using the Akaike Information Criterion (AIC) (Akaike, 1987).

## 3. Results

Results generated by the analysis procedures (illustrated in Figure 1) are presented following the progression from Figure 1A through 1C.

### 3.1. Discrete connectome states have distinct spatial and temporal profiles

As a first step to quantifying dynamic connectome features, we identify discrete connectome states, i.e., whole-brain recurrent connectivity patterns, then show that the states differ from each other in FC strength of specific networks, topological segregation-integration (Modularity), and Fractional Occupancy. To identify the states, we employed a data-driven approach using hidden Markov modeling (HMM) (Vidaurre et al., 2016). To ensure that results are not limited to the specific *a priori* selected number of states (*K*), we applied two different *K* within the range of prior EEG/MEG studies (Baker et al., 2014; Vidaurre et al., 2016; Quinn et al., 2018; Hunyadi et al., 2019; Coquelet et al., 2022). We report outcomes for *K* = 6 and replicate results for *K* = 4 in *Supplementary Information*.

As expected, the states identified by the HMM switched rapidly. The dwell time spent in individual occurrences of states were in the sub-second range for all EEG frequencies (mean across the *K* states and all subjects for delta (414.78 ± 293.0 ms), theta (344.53 ± 329.70 ms), alpha (344.01 ± 417.73 ms), beta (230.67 ± 408.14 ms), and gamma (250.05 ± 736.48 ms)).

Differences in FC strength across connectome states were quantified with a method that accounts for multiple comparisons in graph space (Network-Based Statistics (NBS; Zalesky et al., 2010). Figure 2C illustrates the cluster of connections with cross-state FC differences (NBS FWER-corrected *P^†^* < 0.05) in alpha band, consisting of 70 out of 1431 (5%) connections. Similar observations were made for all other EEG frequency bands (Figure S6 for data-driven set of clusters of connections with different densities). In addition to examining the NBS-derived cluster, we conducted extensive exploratory analyses to assess the involvement of all canonical ICNs. We performed an equivalent one-way ANCOVA of the factor state on FC between each possible pair of canonical ICNs. The results demonstrated that all pairs of ICNs contributed to FC differences across states (*Table S1* for *K*s of 6 and 4 across all frequency bands).

**Figure 2.**
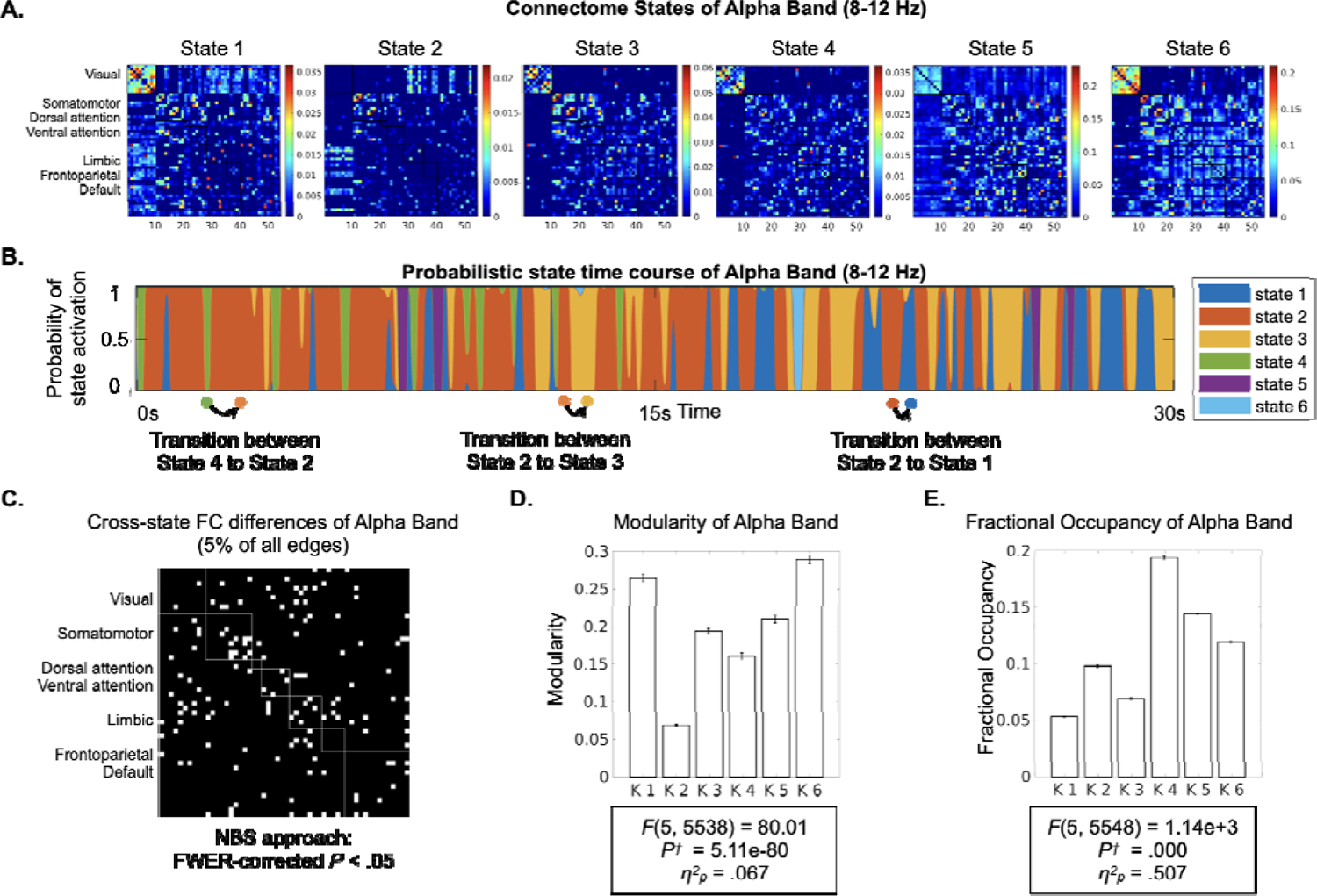
HMM states and state-dissociating features. (A) From band-specific leakage-corrected EEG signals (amplitude envelopes) concatenated over all subjects, HMM estimates connectome states that each have characteristic FC matrix. The FC matrices reflect amplitude coupling among all region pairs. Here, the connectome states’ FC matrices for the alpha band (8-12 Hz) are provided as examples (See full visualization of the states for all canonical bands in *Figure S4*). The rows and columns represent 54 regions organized according to their membershi to canonical intrinsic connectivity networks (ICNs listed on the left) (Yeo et al., 2011). (B) HMM estimates a specific (probabilistic) state time course for each subject indicating when each state is active. An approximately two-minute section of the state time course is visualized for one subject exemplifying periods occupied by each state and the transitions across states. (C) The binary matrix shows data-driven clusters of connections whose FC strength differed significantly across the six states. Specifically, *F* values from a connection-wise ANCOVA of the factor state were threshold at 5% connection density and entered Network-Based Statistics (NBS) to control for multiple comparisons (Zalesky et al., 2010). For the ensuing significant cluster of connections, we provide the (D) Modularity, and (E) Fractional Occupancy for each state. *F* values are reported for one-way ANCOVAs of the factor state (6 levels) for each variable, adjusted for age and sex. Strong differences across states in all three measures suggest distinct spatial and temporal features of each state. *P^†^*: *P* values Bonferroni-corrected for 20 tests (four multivariate features and five frequency bands), η*^2^*: Partial Eta squared effect size.

Moving beyond the specific sets of connections, we investigated cross-state differences in the global topology of the connectome, specifically focusing on Modularity (Newman, 2006; Rubinov and Sporns, 2010) due to its functional relevance (Sadaghiani et al., 2015; Shine and Poldrack, 2018). A one-way ANCOVA of the factor state showed significant difference in Modularity across the connectome states in alpha band (Figure 2D, *F*(_5_, _5538_) = 80.01, *P^†^* = 5.11e-80, η*^2^* = .067). Similarly, this effect was present in all other EEG frequency bands (*Table S2* for *K*s of 6 and 4 across all frequency bands).

Beyond the spatial features, the connectome states showed distinctive temporal characteristics. An equivalent one-way ANCOVA of the factor state on Fractional Occupancy revealed significant differences in the proportion of time spent in each state for the alpha band (Figure 2E, *F*(_5_, _5548_) = 1.14e+03, *P^†^* < 0.001, where *P^†^*is the *P* value Bonferroni-corrected for 20 tests, η*^2^* = .507). Similar observations were made for all other EEG frequency bands (*Table S2* for *K*s of 6 and 4 across all frequency bands).

In summary, these findings confirm that the rapid (sub-second) dynamics of spontaneous connectivity can be characterized as non-random sequences of six discrete connectome states, exhibiting differences in spatial organization, global topology, and proportion of occurrence.

### 3.2. Multivariate temporal features of the dynamic connectome are heritable

We tested the hypothesis that subject pairs with similar genetic makeup had more similar multivariate connectome dynamics features than subjects with less genetic relatedness. Specifically, the multivariate features included Fractional Occupancy (1 × *K*), Transition Probability matrix (*K* × *K*), FC_Time-Varying_ of data-driven clusters (1 × *K*), and Modularity_Time-Varying_ (1 × *K*) (cf. Figure 1A). The similarity of each multivariate feature between a given pair of subjects was quantified as Euclidean distance. Distance values entered a one-way ANCOVA of the factor sibling status with three levels, including monozygotic (MZ) twins, sex-matched dizygotic (DZ) twins, and sex-matched pairs of unrelated individuals, adjusted for age and sex. No subjects overlapped between groups.

We found that temporal features describing the dynamic trajectory of connectome state transitions are heritable in theta, alpha, beta, and gamma bands, but not in delta band (Figure 3). That is, genetically closer subject pairs have more similar Fractional Occupancy and Transition Probability phenotypes, compared to less genetically related pairs: Fractional Occupancy in theta (*F*(*_2_, _458_*) = 9.01*, P^†^* = .003), alpha (*F*(*_2_, _458_*) = 16.80*, P^†^* = 1.82e-06), beta (*F*(*_2_, _458_*) = 9.95*, P^†^* = .001), and gamma (*F*(*_2_, _458_*) = 11.17, *P^†^* = 3.65e-04) and Transition Probability in theta (*F*(*_2_, _458_*) = 7.55, *P^†^* = .012), alpha (*F*(*_2_, _458_*) = 14.51, *P^†^* = 1.55e-05), and gamma (*F*(*_2_, _458_*) =, *P^†^* = 2.97e-05). This impact of sibling status on temporal features was consistently large and independent of the chosen number of states (cf. Figure 3 and *Figure S5*). Note that there was no effect of sibling status in surrogate data lacking time-varying dynamics but with preserved static covariance structure (cf. Null Model section 2.6): Fractional Occupancy in delta through gamma (*F*_(2, 458)_ = 2.99, *F*_(2, 458)_ = 2.12, *F*_(2, 458)_ = .55, *F*_(2, 458)_ = .06, and *F*_(2, 458)_ = .07 respectively, all *P*^†^ = 1.00) and Transition Probability in delta through gamma (*F*_(2, 458)_ = 2.89, *F*_(2, 458)_ = .39, *F*_(2, 458)_ = .38, *F*_(2, 458)_ = 1.08, and *F*_(2, 458)_ = .54, all *P*^†^ = 1.00).

**Figure 3.**
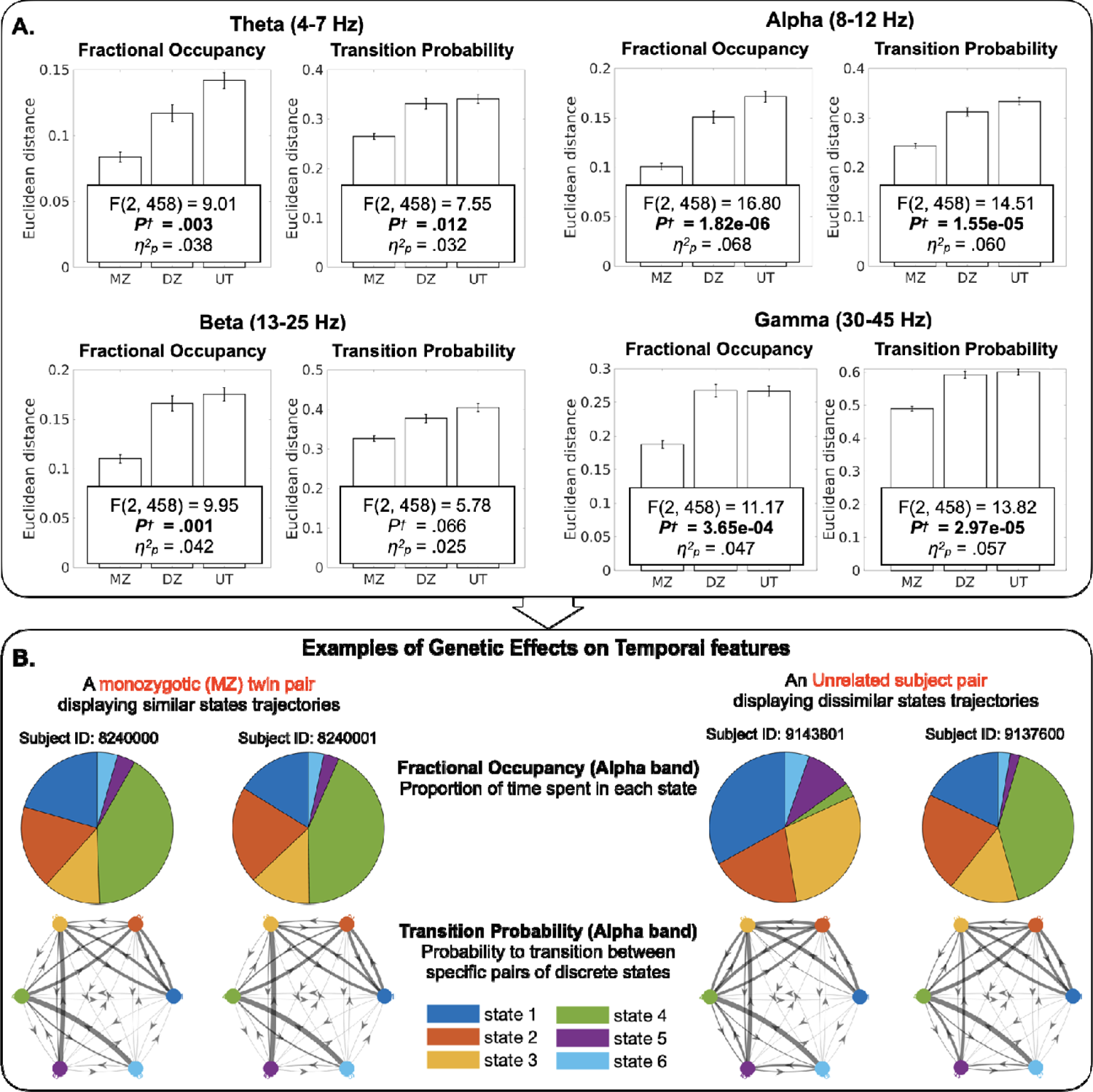
Heritability of temporal features of the dynamic connectome. **(A)** Heritability of each of the multivariate connectome dynamics features was assessed separately by one-way ANCOVAs of the factor sibling status (thre levels: monozygotic twins (MZ), sex-matched dizygotic twins (DZ), and sex-matched pairs of unrelated individuals), adjusted for age and sex. The main effect of sibling status indicates the heritability, or genetic effect. Bar graphs show that genetically more similar subject pairs have shorter Euclidean distance, indicating higher similarity of a give temporal feature. This effect of sibling status was large and independent of the chosen number of states (see *Figure S5*). *P^†^*: P values Bonferroni-corrected for 20 tests (four multivariate features and five frequency bands), η*^2^*: Partial Eta squared effect size. **(B)** A visual illustration of the effects exemplified for two subject pairs, one from the MZ grou (left) and the other from the unrelated subject group (right). While the MZ twins display high similarity of Fractional Occupancy and Transition Probabilities, both temporal features are dissimilar across the unrelated pair.

Contrary to the *temporal* features, we did not find support that was robust (i.e., invariant to methodological choices) for heritability of the *spatial* features, describing how connectome states are spatially instantiated in individuals. Specifically, outcomes of equivalent ANCOVAs for Modularity_Time-Varying_ and FC_Time-Varying_ of data-driven clusters showed no impact of sibling status, irrespective of frequency bands and chosen number of states (*Table S3*). This lack of robust heritability was confirmed by a subsequent variance-component genetic analysis (see **Results** III). Our sample size permitted detecting, at 80% power, effects of small size (η^2^ = .021, equivalent to *f* = .145, or larger). Therefore, if for spatial features heritability produced effects smaller than the detectable size, these effects would be of low practical impact. Further, we provide additional Bayesian Factor values to directly assess the probability of H_0_ (i.e., the null hypothesis that there is no effect of sibling status) against H_1_ (*Table S3*). Indeed, the Bayes Factor for Modularity_Time-Varying_ and FC_Time-Varying_ of data-driven clusters showed that the data are more likely to occur under H_0_ than under H_1_. For example, Modularity_Time-Varying_ in alpha band showed anecdotal support for the null hypothesis (BF_01_ = 1.72), whereas FC_Time-Varying_ of data-driven clusters in alpha band presented strong support for the null hypothesis (BF_01_ = 33.57; suggesting that data are 34 times more likely to occur under H_0_ than under H_1_). Notably, the lack of evidence for the heritability of spatial connectome dynamics features aligns with an equivalent null result in our preceding fMRI study (Jun et al., 2022).

To ensure that the absence of a robust outcome for spatial features was not influenced by a narrow feature selection, we performed additional equivalent analyses on exploratory spatial features: FC_Time-Varying_ of data-driven clusters defined across four other connection densities (1∼4% across different number of states; *Table S3*) and FC_Time-Varying_ of all 21 possible pairs among the seven ICNs (*Tables S4*). Consistent with our findings for the main spatial features, ANCOVAs of the factor sibling status and Bayes Factors of all of the exploratory spatial features provided anecdotal to decisive evidence for H_0_, i.e., the lack of heritability. These findings strongly contrast the observations for temporal features, where Bayes Factor of both Fractional Occupancy and Transition Probability showed that the data are significantly more likely to occur under H_1_ than under H_0_: Fractional Occupancy in theta (BF_10_ = 1.07e+02), alpha (BF_10_ = 1.77e+05), beta (BF_10_ = 2.99e+02), gamma (BF_10_ = 9.51e+02) and Transition Probability in theta (BF_10_ = 26.5), alpha (BF_10_ = 2.08e+04), beta (BF_10_ = 5.13), gamma (BF_10_ = 9.72e+03).

Importantly, the effect sizes of significant multivariate Fractional Occupancy and Transition Probability were notably larger (with mean η*^2^* of .049 across all frequency bands; Figure 3) compared to those of the *individual* (i.e., state-by-state) components of the multivariate features (with mean η*^2^* of .016 across all frequency bands; *Table S5*). Therefore, our findings demonstrate that the dynamic trajectory of rapid EEG connectome state transitions are robustly heritable, predominantly when considered as multivariate patterns, rather than as individual state-specific components.

### 3.3. Genetic effects account for substantial variability in temporal connectome dynamics

We then quantified the extent to which genetic variance contributes to phenotypic variance using structural equation modeling, a technique commonly employed in classical twin studies (Falconer, 1990). While the model is traditionally applied to univariate phenotypes, the above-described one-way ANCOVAs of the factor sibling status suggest that connectome dynamics features are inherited predominantly as *multivariate* patterns. Therefore, we adapted the model to accommodate multivariate phenotypes by quantifying the subject-wise Euclidean distance of multivariate features from a “null” point of origin (from dynamics-free surrogate data; Figure 1 and **Materials and Methods 2.6 Null models**).

A substantial portion of phenotypic variance in the temporal features was explained by genetic variance in the genetic models, adjusted for age and sex (Table 1; *Table S6* for *K* = 4). Specifically, Transition Probability in alpha, beta, and gamma bands, as well as Fractional Occupancy in beta and gamma bands were explained by the ADE model. In the ADE model, A (additive genetic effect) and D (dominant genetic effect) together estimate broad-sense heritability (*H^2^*). Conversely, Transition Probability in the theta band was described by the ACE model, where narrow-sense heritability (*h^2^*) was estimated using the A variance component. Notably, we found substantial heritability of Transition Probability (*h^2^* of the theta band = 38.4% [.03, .74], *H^2^* of the alpha band = 63.3% [.56, .71], *H^2^* of the beta band = 22.6% [.08, .37], and *H^2^* of the gamma band = 40% [.28, .52]) and Fractional Occupancy (*H^2^* of the beta band = 44.3% [.33, .55] and *H^2^* of the gamma band = 39.8% [.28, .52]). In all cases, the fitness of the nested models (i.e., AE, CE, or DE) was not significantly better than the ACE or ADE model. These outcomes indicate that genetics contribute substantially to the temporal features of connectome dynamics.

**Table 1.**
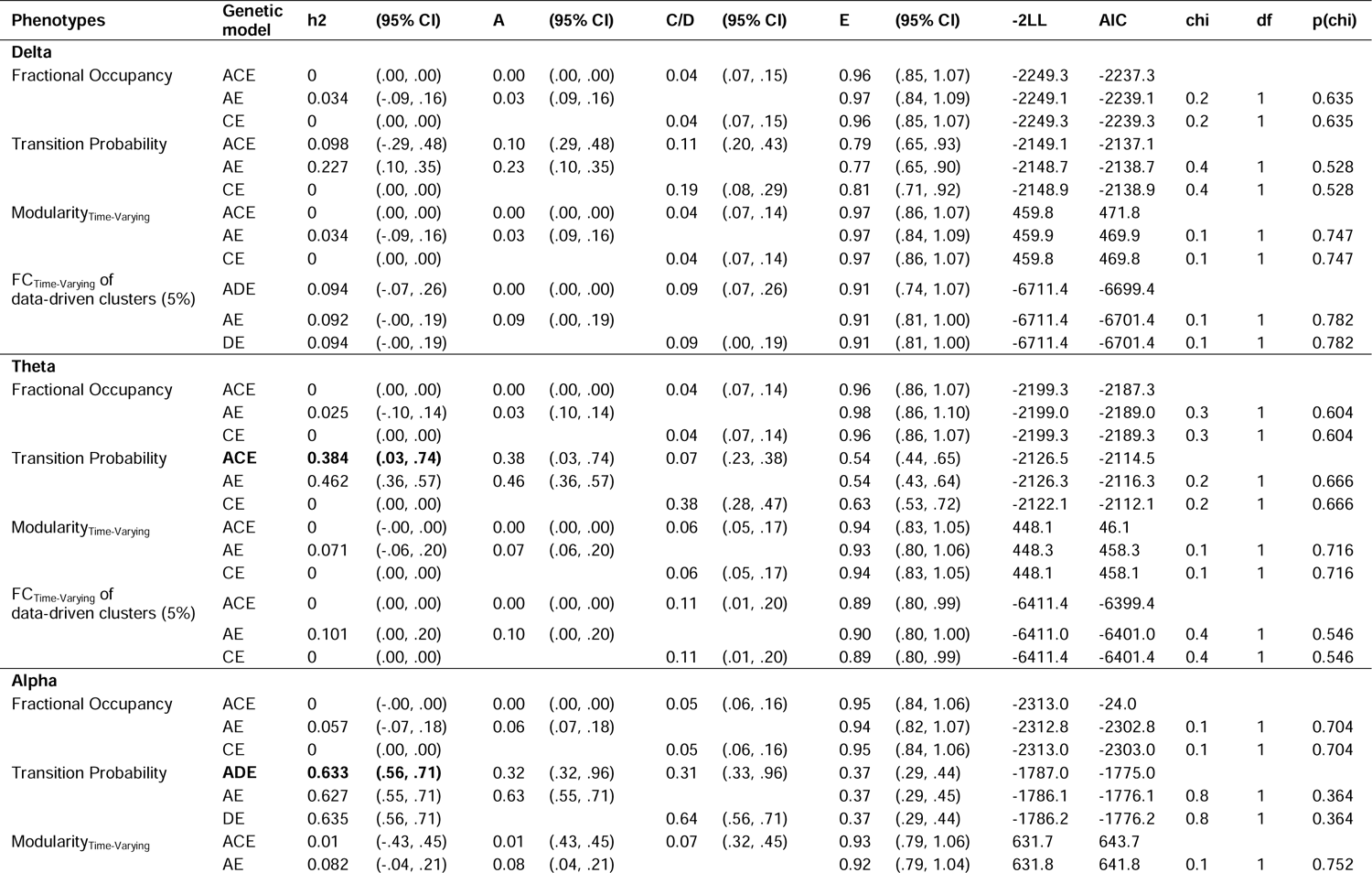
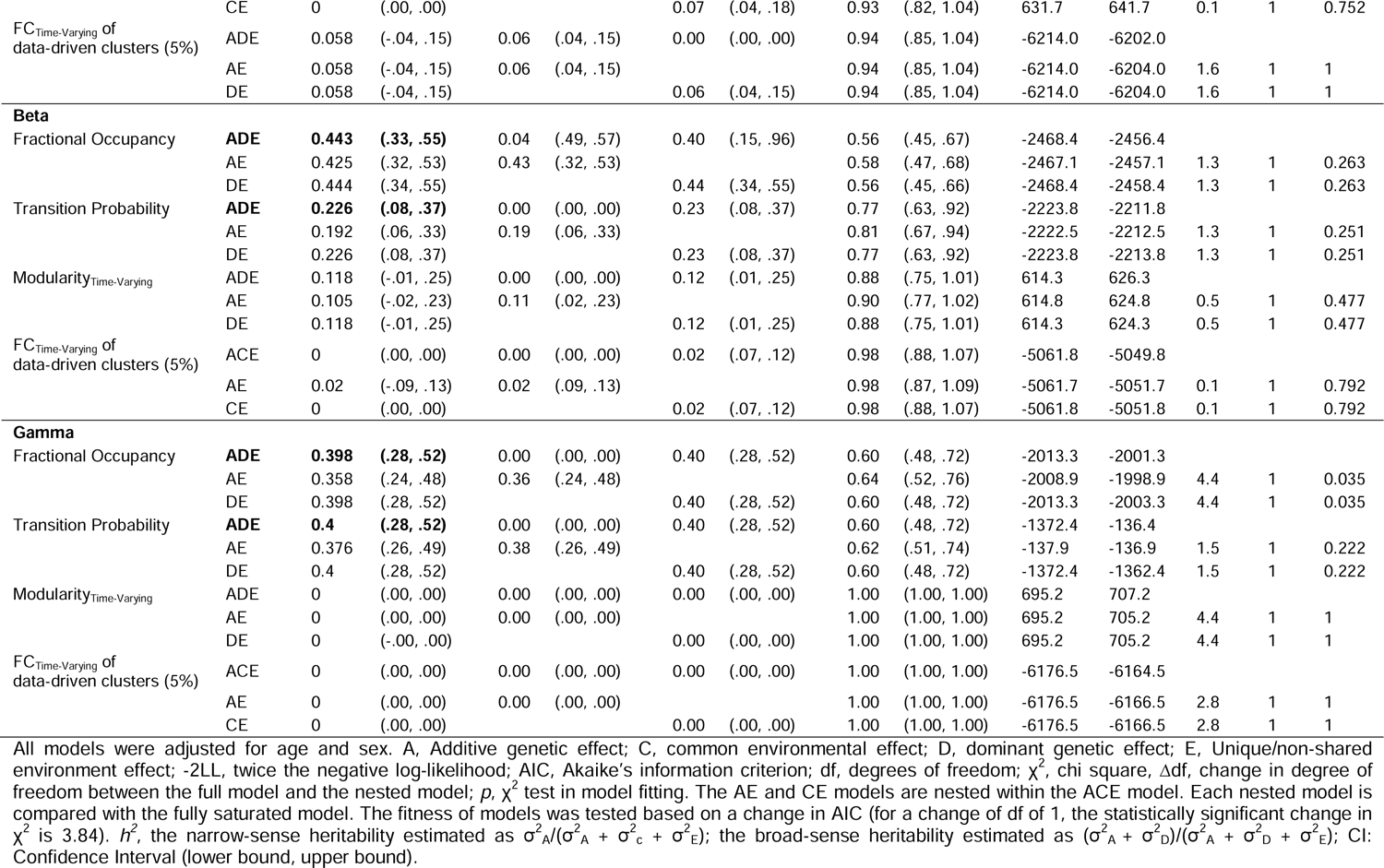
Variance-component model parameter estimates of the dynamic connectome features (K = 6).

Consistent with the ANCOVA-based heritability findings (*Tables S3-4*), the genetic models did not support genetic effects on either spatial feature (Modularity_Time-Varying_ and FC_Time-Varying_ of data-driven clusters; Table 1 and *Table S6* for *K* of 4). Specifically, the heritability (*h^2^*) of spatial features was estimated as zero or its 95% confidence interval crossed zero, supporting the null hypothesis, i.e., lack of heritability.

## Discussion

The use of source-space electrophysiological signals allowed us to investigate rapidly emerging and dissolving patterns of spatially localized connectivity networks and their transitions at the whole-brain level. Bringing this approach to a large cohort, we established genetic effects on rapid connectome dynamics in specific frequency bands. Overcoming the limited temporal resolution of prior heritability investigations of slow connectome dynamics in fMRI (cf. infra-slow (< 0.1 Hz) fMRI BOLD signal; (Vidaurre et al., 2017; Barber et al., 2021; Jun et al., 2022), the current findings shed light on sub-second timescales highly relevant to cognitive processes. As an additional innovation, we embraced the multi-dimensional nature of dynamic connectome features (Jun et al., 2022), as these features collectively encompass patterns from multiple connectome states. Reinforcing the multi-dimensional view, the heritability effect size was larger for *multivariate* features (Figure 3 and *Figure S5*) than for state-wise scalar features (cf. *Table S5*). Quantitative modeling with the multivariate features showed that the genetic influence on the rapid sequencing of connectome states was substantial (22-63% of variance explained).

Previous twin studies have found considerable genetic effects on multiple EEG features, however predominantly focusing on stable rather than time-varying characteristics. Specifically, in terms of the power spectrum, heritability was particularly evident in individuals’ alpha peak frequency (71% to 83% phenotypic variance explained (Posthuma et al., 2001; Smit et al., 2006) and alpha power (79% to 93% variance explained, depending on scalp location and age of cohort (Smit et al., 2005, 2006)). The degree of heritability observed beyond the alpha band has been somewhat less pronounced; in direct comparisons across the spectrum the highest heritability of spectral power was observed at the alpha peak frequency, while it was lower in theta and delta bands (Smit et al., 2005).

Besides the EEG power spectrum, static (time-averaged) M/EEG FC measures were found to be subject-specific (cf. fingerprinting; (Kong et al., 2019; Kabbara et al., 2021; Sareen et al., 2021)) and under genetic influence. For example, the heritability estimates of static EEG FC ranged from 27% to 75%, primarily observed in alpha and beta bands (Posthuma et al., 2005; Schutte et al., 2013). Additionally, heritability estimates for graph theoretical measures of the static EEG FC matrix ranged from 46% to 89% for clustering coefficients and from 37% to 62% for average path length (Smit et al., 2008). However, it is important to note that the above-mentioned studies were performed in sensor-space and employed synchronization likelihood for network estimation, a measure that may be susceptible to volume conduction artifacts. A *source-space* MEG heritability study correcting such artifacts estimated the heritability of amplitude coupling in individual edges, which averaged across edges reached 8% for the alpha band and 19% for the beta band (Colclough et al., 2017).

Our study significantly extends this important previous research by establishing heritability of rapid whole-brain connectome dynamics, derived from source-localized (and leakage-corrected) EEG. Our quantitative heritability estimates fall within the range reported in the above-described studies that focused on stable EEG spectral properties and static FC investigations. The strongest genetic influence in our study was observed in the alpha band, paralleling prior heritability estimates of the power spectrum. While this dominance may result from the high signal-to-noise ratio of the alpha band, the strong alpha signal in itself likely reflects an important functional role of this frequency in cognition (Palva and Palva, 2011; Klimesch, 2012; Sadaghiani and Kleinschmidt, 2016). In general, and for all frequency bands, it is the rapid changes that are thought to be particularly critical for cognitive processes, which are inherently dynamic at numerous timescales (Gratton, 2018). By addressing electrophysiological processes from a time-varying perspective, the current study establishes the heritability of cognitively significant rapid brain state changes. Our separate study in the same cohort directly confirms the predicted implications of such rapid connectome dynamics for individual differences in cognitive abilities (Jun et al., 2024; Submitted).

Notably, robust evidence for a genetic influence was found for Transition Probabilities in all bands except delta but was observed for Fractional Occupancy in the higher bands only (beta and gamma). While Transition Probability and Fractional Occupancy are not fully independent measures, they contain non-overlapping information about connectome dynamics. For example, a state with particularly high Fractional Occupancy is likely to have high values as initial state and target state in the Transition Probability matrix. Despite such dependence, however, two hypothetical subjects with highly comparable Fractional Occupancy values across the *k* states may still have substantially different sequencing, and thus transition probabilities, across the states. This sequencing, as suggested by our findings, is under strong genetic influence broadly across electrophysiological timescales. At least at infra-slow timescales (typically observed in fMRI), such brain state changes are in part driven by the spatially broad but structured influence of ascending modulatory neurotransmitter systems (Klaassens et al., 2017; Shine et al., 2018; Lord et al., 2019). Numerous genetic polymorphisms with functional impact are known within receptors, transporters, and enzymes of these systems (Kautzky et al., 2015; Sadaghiani et al., 2017; de Rojas et al., 2021). Future causal (e.g., neuropharmacological interventions and subcortical micro-stimulation) or modelling studies could assess to what degree similar neuromodulatory process are at play in the individually specific connectome dynamics at rapid timescales.

Consistent with our previous fMRI investigation (Jun et al., 2022), the results concerning the spatial features of connectome states largely supported the null hypothesis of the absence of heritability for a wide range of features. Specifically, when examining our primary spatial features, i.e., cluster-based FC_Time-Varying_ and Modularity_Time-Varying_, there was no discernible influence of sibling status on the phenotypic similarity between subjects across different number of states. This lack of heritability for cluster-based FC_Time-Varying_ was further corroborated by exploring additional connection densities (*Table S3*). Further, in an exhaustive exploratory assessment of 21 ICN pairs, FC_Time-Varying_ exhibited no observable influence from sibling status under different methodological alternatives (*Table S4*). Therefore, our study suggests that genetic effects primarily contribute to how the connectome *transitions* across different states, rather than the precise way in which the states are spatially instantiated in individuals. It is remarkable that this dissociation between temporal and spatial features of connectome dynamics holds across the full spectrum of connectivity timescales, from infra-slow (fMRI) through gamma band (EEG). Still, it is important to acknowledge that factors beyond genetics, such as individuals’ experiences and learning, play a substantial role in shaping subject-specific connectomes and their spatial patterns, as indicated by the significant contribution of common and random environmental variances to the spatial features in Table 1.

Our study is subject to several limitations and methodological considerations. While we provide results separately for each canonical frequency band, this approach does not assume or necessitate the bands to be discretely separable or oscillatory in nature. The approach is equally compatible with a more general view that the bands represent electrophysiological processes at different speed within a larger 1/f spectrum. Further, we defined the boundaries of the frequency bands according to common conventions in the field rather than according to the individual’s power spectrum. Because the latter (especially the alpha peak frequency commonly used to anchor individual bands) is highly heritable (see above), defining the bands individually may strengthen the observed heritability of connectome dynamics. However, a non-individualized definition of bands is unlikely to result in false positives in terms of such heritability. Another consideration is that while the set of spatial features of connectome dynamics in our main and supplementary reports was large, it is necessarily inexhaustive. Other dynamic spatial features not explored in this study could potentially exhibit heritability and call for future investigation.

In conclusion, our findings provide the first evidence of genetic influence on rapid transitions between whole-brain source-space EEG connectome states and the proportion of time spent in each state. In combination with our previous findings in fMRI-derived dynamics (Jun et al., 2022), the evidence of a genetic basis of connectome state trajectories extends the full breadth of connectivity timescales from infra-slow to gamma band. Extensive prior work has established that various aspects of brain anatomy are heritable, including how the brain is wired in terms white matter tracts (Kochunov et al., 2015; Sha et al., 2023). The genetic influence on wiring has further been extended to the time-averaged functional connectome (Schutte et al., 2013; Sinclair et al., 2015; Colclough et al., 2017), reflecting which brain regions have a strong (or respectively weak) average tendency to coordinate their activity. While this important prior research has focused on the brain’s stable structural/functional architecture, the current work shifted the focus to the brain’s dynamic behavior (i.e., what the brain *does* in terms of time-varying dynamics). Interestingly, the shift to time-varying connectome states showed that genetics impacts the *act of transitioning* across states rather than the spatial organization of those states (at least among the features we studied). This observation gains particular importance in light of the central role of rapid electrophysiological dynamics in brain function and cognition (Gratton, 2018). Our findings may inform the identification of functionally relevant genetic polymorphisms and the development of connectome-based biomarkers at timescales particularly relevant to cognitive processes.

## Acknowledgements

We thank Dr. Andre Altmann for his extensive guidance in analytic approaches, and Drs. Jonathan Wirsich and Thomas Alderson for guidance in data preprocessing. Computational resources for this work were provided by the Minnesota Supercomputing Institute at the University of Minnesota Informatics Institute. The Center for Magnetic Resonance Research (supported by Grant Nos. NIBIB P41 EB027061 and 1S10OD017974-01) at the University of Minnesota provided resources that contributed to the MRI-related results reported within this article. The original data collection of the data analyzed in this paper was funded by NIH grants R37 DA05147 and R01 DA036216. This work was partly supported by the National Institute for Mental Health (1R01MH116226 to Sepideh Sadaghiani).

