## Supplementary File for "Rapid dynamics of electrophysiological connectome states are heritable"

**Supplementary Information**

**SI Result. Alternative multivariate genetic variance component model.**

Our previous work (Jun et al., 2022) provided a novel approach to adjust the common structural equation model traditionally used in classical twin studies of univariate phenotypes (Falconer, 1990) to fit multivariate phenotypes. This adjustment was made available by using the similarity (or Euclidean distance) between subject’s multivariate feature and a given origin point (**Materials and Methods** **– 2.9. Similarity estimation and heritability testing**; Fig. 1C). To provide further support for the validity of the adaptation to multivariate phenotypes, we applied an alternative model (Ge et al., 2016) to estimate the heritability of multivariate features. This alternative weighted sum approach computes heritability of the multivariate phenotype as the cumulative heritability of individual components (cf. *Tables S8-9* for formula). However, it should be noted that this cumulative approach will overestimate heritability for highly collinear phenotype features (since the shared variance explained by collinear features enters the sum multiple times), and as such is not optimal for Fractional Occupancy and Transition Probability.

Despite this conceptual difference to our multivariate model, the cumulative heritability model resulted in comparable outcomes. The sum of weighted heritability for Fractional Occupancy in the delta, theta, alpha, beta, and gamma bands were 14.9%, 22.2%, 47.6%, 38.5%, and 23.5%, respectively (*Tables S8*). Similarly, the sum of weighted heritability for Transition Probability in the delta, theta, alpha, beta, and gamma bands were 10.5%, 22.2%, 37.7%, 22.1%, and 21.7%, respectively (*Tables S9*). These values are reasonably close to the heritability (near or within the confidence intervals) estimated in our multivariate approach (Table 1). Specifically, consistent with our multivariate approach, the values support highest heritability in the alpha band. The converging results support the validity of our multidimensional space-based approach, including its origin point determined from surrogate data.

**Supplementary Tables**

### Table S1. ANCOVAs of the factor connectome state (*K* levels) on FC_Time-Varying_ of all Intrinsic connectivity network (ICN) pairs.

| **Variables** | ***K* = 4** |  |  | ***K* = 6** |  |  |
| --- | --- | --- | --- | --- | --- | --- |
|  | ***F* states** | ***P*^†^** | ***η^2^_p_*** | ***F* states** | ***P*^†^** | ***η^2^_p_*** |
| **Delta band** |  |  |  |  |  |  |
| VIS-SMN_Time-Varying_ | 8.19E+02 | 0 | 0.399 | 1.98E+02 | 2.55E-192 | 0.151 |
| VIS-DAN_Time-Varying_ | 8.62E+01 | 6.84E-52 | 0.065 | 2.60E+02 | 3.14E-248 | 0.190 |
| VIS-VAN_Time-Varying_ | 1.12E+03 | 0 | 0.475 | 4.15E+02 | 0 | 0.272 |
| VIS-Limbic_Time-Varying_ | 8.96E+02 | 0 | 0.421 | 1.59E+02 | 9.08E-157 | 0.126 |
| VIS-FPN_Time-Varying_ | 2.93E+02 | 9.40E-169 | 0.192 | 5.42E+01 | 4.50E-53 | 0.047 |
| VIS-DMN_Time-Varying_ | 5.14E+02 | 2.85E-277 | 0.295 | 1.73E+02 | 2.50E-169 | 0.135 |
| SMN-DAN_Time-Varying_ | 6.37E+02 | 0 | 0.341 | 5.52E+01 | 3.72E-54 | 0.047 |
| SMN-VAN_Time-Varying_ | 3.01E+02 | 6.06E-173 | 0.197 | 6.46E+01 | 1.04E-63 | 0.055 |
| SMN-Limbic_Time-Varying_ | 6.41E+02 | 0 | 0.342 | 6.40E+01 | 4.30E-63 | 0.055 |
| SMN-FPN_Time-Varying_ | 7.39E+02 | 0 | 0.375 | 8.93E+01 | 1.40E-88 | 0.074 |
| SMN-DMN_Time-Varying_ | 1.14E+03 | 0 | 0.480 | 2.13E+02 | 1.89E-206 | 0.161 |
| DAN-VAN_Time-Varying_ | 1.07E+02 | 2.51E-64 | 0.080 | 4.59E+02 | 0 | 0.293 |
| DAN-Limbic_Time-Varying_ | 1.88E+02 | 1.82E-111 | 0.132 | 4.16E+01 | 3.74E-40 | 0.036 |
| DAN-FPN_Time-Varying_ | 4.51E+02 | 1.28E-247 | 0.268 | 1.09E+02 | 1.20E-107 | 0.089 |
| DAN-DMN_Time-Varying_ | 8.18E+02 | 0 | 0.399 | 2.18E+02 | 2.35E-211 | 0.165 |
| VAN-Limbic_Time-Varying_ | 5.63E+02 | 1.80E-299 | 0.314 | 8.46E+01 | 6.15E-84 | 0.071 |
| VAN-FPN_Time-Varying_ | 8.52E+01 | 2.61E-51 | 0.065 | 6.68E+01 | 6.15E-66 | 0.057 |
| VAN-DMN_Time-Varying_ | 9.32E+01 | 3.83E-56 | 0.070 | 2.04E+02 | 6.00E-198 | 0.155 |
| Limbic-FPN_Time-Varying_ | 8.45E+02 | 0 | 0.407 | 1.04E+02 | 2.03E-103 | 0.086 |
| Limbic-DMN_Time-Varying_ | 9.45E+02 | 0 | 0.434 | 1.12E+02 | 4.97E-111 | 0.092 |
| FPN-DMN_Time-Varying_ | 1.57E+03 | 0 | 0.561 | 4.94E+02 | 0 | 0.308 |
| **Theta band** |  |  |  |  |  |  |
| VIS-SMN_Time-Varying_ | 8.37E+02 | 0 | 0.405 | 8.34E+01 | 1.12E-82 | 0.070 |
| VIS-DAN_Time-Varying_ | 2.36E+00 | 7.31E+00 | 0.002 | 3.23E+01 | 1.47E-30 | 0.028 |
| VIS-VAN_Time-Varying_ | 1.24E+03 | 0 | 0.502 | 3.57E+02 | 0 | 0.244 |
| VIS-Limbic_Time-Varying_ | 9.70E+02 | 0 | 0.441 | 1.53E+02 | 5.77E-151 | 0.122 |
| VIS-FPN_Time-Varying_ | 3.29E+02 | 3.82E-187 | 0.211 | 3.35E+01 | 9.63E-32 | 0.029 |
| VIS-DMN_Time-Varying_ | 5.52E+02 | 3.74E-294 | 0.310 | 7.12E+01 | 2.29E-70 | 0.060 |
| SMN-DAN_Time-Varying_ | 3.58E+02 | 4.19E-202 | 0.226 | 1.49E+02 | 5.67E-147 | 0.119 |
| SMN-VAN_Time-Varying_ | 3.58E+02 | 4.15E-202 | 0.226 | 5.62E+01 | 3.88E-55 | 0.048 |
| SMN-Limbic_Time-Varying_ | 7.16E+02 | 0 | 0.368 | 5.58E+01 | 8.53E-55 | 0.048 |
| SMN-FPN_Time-Varying_ | 9.89E+02 | 0 | 0.446 | 1.69E+02 | 9.89E-166 | 0.132 |
| SMN-DMN_Time-Varying_ | 1.05E+03 | 0 | 0.461 | 2.09E+02 | 2.14E-202 | 0.159 |
| DAN-VAN_Time-Varying_ | 3.91E+01 | 7.70E-23 | 0.031 | 8.20E+01 | 2.99E-81 | 0.069 |
| DAN-Limbic_Time-Varying_ | 7.65E+01 | 5.72E-46 | 0.059 | 1.97E+01 | 1.54E-17 | 0.018 |
| DAN-FPN_Time-Varying_ | 2.24E+02 | 1.14E-131 | 0.154 | 6.18E+01 | 7.18E-61 | 0.053 |
| DAN-DMN_Time-Varying_ | 3.90E+02 | 7.51E-218 | 0.241 | 1.08E+02 | 1.19E-106 | 0.089 |
| VAN-Limbic_Time-Varying_ | 5.68E+02 | 2.11E-301 | 0.316 | 1.20E+02 | 1.28E-118 | 0.098 |
| VAN-FPN_Time-Varying_ | 8.86E+01 | 2.54E-53 | 0.067 | 3.31E+01 | 2.01E-31 | 0.029 |
| VAN-DMN_Time-Varying_ | 1.07E+02 | 1.06E-64 | 0.080 | 4.50E+01 | 1.27E-43 | 0.039 |
| Limbic-FPN_Time-Varying_ | 6.17E+02 | 0 | 0.334 | 5.21E+01 | 6.32E-51 | 0.045 |
| Limbic-DMN | 8.96E+02 | 0 | 0.421 | 1.12E+02 | 2.25E-111 | 0.092 |
| FPN-DMN | 9.14E+02 | 0 | 0.426 | 2.71E+02 | 7.07E-258 | 0.197 |
| **Alpha band** |  |  |  |  |  |  |
| VIS-SMN | 9.45E+01 | 6.69E-57 | 0.071 | 6.42E+02 | 0 | 0.367 |
| VIS-DAN | 2.24E+02 | 2.79E-131 | 0.154 | 1.05E+02 | 2.71E-104 | 0.087 |
| VIS-VAN | 6.35E+02 | 0 | 0.340 | 4.99E+02 | 0 | 0.311 |
| VIS-Limbic | 8.22E+01 | 1.81E-49 | 0.063 | 4.61E+02 | 0 | 0.294 |
| VIS-FPN | 1.48E+01 | 1.44E-07 | 0.012 | 3.49E+02 | 0 | 0.240 |
| VIS-DMN | 5.64E+01 | 1.33E-33 | 0.044 | 6.75E+02 | 0 | 0.379 |
| SMN-DAN | 3.78E+02 | 3.72E-212 | 0.235 | 2.29E+02 | 9.55E-221 | 0.171 |
| SMN-VAN | 5.86E+01 | 5.40E-35 | 0.045 | 4.26E+02 | 0 | 0.278 |
| SMN-Limbic | 1.72E+02 | 1.82E-102 | 0.123 | 6.30E+02 | 0 | 0.363 |
| SMN-FPN | 1.98E+02 | 4.49E-117 | 0.139 | 5.37E+02 | 0 | 0.327 |
| SMN-DMN | 3.03E+02 | 1.17E-173 | 0.197 | 8.00E+02 | 0 | 0.419 |
| DAN-VAN | 4.61E+02 | 1.80E-252 | 0.273 | 9.53E+01 | 1.34E-94 | 0.079 |
| DAN-Limbic | 1.84E+02 | 6.45E-109 | 0.130 | 1.04E+02 | 4.92E-103 | 0.086 |
| DAN-FPN | 2.94E+02 | 2.32E-169 | 0.193 | 1.60E+02 | 7.36E-157 | 0.126 |
| DAN-DMN | 4.44E+02 | 4.06E-244 | 0.265 | 2.98E+02 | 2.80E-281 | 0.212 |
| VAN-Limbic | 4.10E+01 | 5.21E-24 | 0.032 | 4.15E+02 | 0 | 0.272 |
| VAN-FPN | 8.46E+01 | 6.40E-51 | 0.064 | 2.31E+02 | 2.82E-222 | 0.172 |
| VAN-DMN | 6.97E+01 | 8.06E-42 | 0.054 | 4.06E+02 | 0 | 0.268 |
| Limbic-FPN | 1.95E+02 | 2.16E-115 | 0.137 | 3.94E+02 | 0 | 0.262 |
| Limbic-DMN | 2.02E+02 | 3.02E-119 | 0.141 | 7.10E+02 | 0 | 0.391 |
| FPN-DMN | 4.28E+02 | 1.83E-236 | 0.258 | 6.60E+02 | 0 | 0.373 |
| **Beta band** |  |  |  |  |  |  |
| VIS-SMN | 4.51E+02 | 1.39E-247 | 0.268 | 2.43E+02 | 2.19E-228 | 0.211 |
| VIS-DAN | 1.20E+02 | 3.13E-72 | 0.089 | 8.33E+01 | 6.01E-82 | 0.084 |
| VIS-VAN | 7.26E+02 | 0 | 0.371 | 2.71E+02 | 4.67E-252 | 0.230 |
| VIS-Limbic | 3.34E+02 | 5.78E-190 | 0.214 | 2.50E+02 | 7.00E-235 | 0.216 |
| VIS-FPN | 4.44E+01 | 3.56E-26 | 0.035 | 8.69E+01 | 1.70E-85 | 0.087 |
| VIS-DMN | 1.52E+02 | 5.16E-91 | 0.110 | 2.36E+02 | 9.32E-223 | 0.206 |
| SMN-DAN | 3.51E+02 | 1.62E-198 | 0.222 | n/a | n/a | n/a |
| SMN-VAN | 1.49E+02 | 3.95E-89 | 0.108 | 1.14E+02 | 7.33E-112 | 0.112 |
| SMN-Limbic | 2.79E+02 | 3.68E-161 | 0.185 | n/a | n/a | n/a |
| SMN-FPN | 4.80E+02 | 2.43E-261 | 0.281 | n/a | n/a | n/a |
| SMN-DMN | 6.24E+02 | 0 | 0.337 | 5.06E+02 | 0 | 0.358 |
| DAN-VAN | 1.84E+02 | 5.36E-109 | 0.130 | n/a | n/a | n/a |
| DAN-Limbic | 7.53E+01 | 2.87E-45 | 0.058 | n/a | n/a | n/a |
| DAN-FPN | 2.93E+02 | 1.66E-168 | 0.192 | n/a | n/a | n/a |
| DAN-DMN | 4.70E+02 | 8.22E-257 | 0.277 | 3.12E+02 | 7.98E-286 | 0.256 |
| VAN-Limbic | 2.27E+02 | 4.74E-133 | 0.156 | 1.98E+02 | 3.52E-189 | 0.179 |
| VAN-FPN | 1.64E+01 | 1.43E-08 | 0.013 | 3.76E+01 | 7.03E-36 | 0.040 |
| VAN-DMN | 4.70E+01 | 8.94E-28 | 0.037 | 9.89E+01 | 3.32E-97 | 0.098 |
| Limbic-FPN | 3.87E+02 | 2.12E-216 | 0.239 | 1.68E+02 | 6.23E-162 | 0.156 |
| Limbic-DMN | 3.77E+02 | 8.99E-212 | 0.235 | 3.02E+02 | 5.66E-278 | 0.250 |
| FPN-DMN | 9.01E+02 | 0 | 0.423 | 3.77E+02 | 0 | 0.293 |
| **Gamma band** |  |  |  |  |  |  |
| VIS-SMN | 4.85E+02 | 9.78E-264 | 0.283 | 2.87E+02 | 1.57E-270 | 0.212 |
| VIS-DAN | 4.55E+01 | 7.36E-27 | 0.036 | 3.25E+01 | 8.82E-31 | 0.030 |
| VIS-VAN | 6.04E+02 | 3.19E-317 | 0.330 | 3.41E+02 | 9.12e-316 | 0.242 |
| VIS-Limbic | 4.62E+02 | 6.87E-253 | 0.273 | 2.77E+02 | 3.06E-262 | 0.206 |
| VIS-FPN | 1.86E+02 | 2.90E-110 | 0.131 | 8.55E+01 | 1.09E-84 | 0.074 |
| VIS-DMN | 2.45E+02 | 6.52E-143 | 0.166 | 1.15E+02 | 1.33E-113 | 0.097 |
| SMN-DAN | 6.51E+02 | 0 | 0.347 | 3.20E+02 | 2.07E-298 | 0.231 |
| SMN-VAN | 2.08E+02 | 8.50E-123 | 0.145 | 1.17E+02 | 4.60E-116 | 0.099 |
| SMN-Limbic | 6.05E+02 | 1.35e-317 | 0.330 | 3.23E+02 | 1.08E-300 | 0.232 |
| SMN-FPN | 5.99E+02 | 5.68e-315 | 0.328 | n/a | n/a | n/a |
| SMN-DMN | 7.37E+02 | 0 | 0.375 | 5.13E+02 | 0 | 0.324 |
| DAN-VAN | 7.22E+01 | 2.48E-43 | 0.055 | 2.03E+01 | 4.26E-18 | 0.019 |
| DAN-Limbic | 2.05E+02 | 4.44E-121 | 0.143 | 1.04E+02 | 9.55E-103 | 0.089 |
| DAN-FPN | 4.79E+02 | 1.27E-260 | 0.280 | n/a | n/a | n/a |
| DAN-DMN | 8.87E+02 | 0 | 0.419 | 4.65E+02 | 0 | 0.303 |
| VAN-Limbic | 3.71E+02 | 1.05E-208 | 0.232 | 2.24E+02 | 9.24E-216 | 0.174 |
| VAN-FPN | 8.48E+01 | 5.12E-51 | 0.065 | 2.28E+01 | 1.06E-20 | 0.021 |
| VAN-DMN | 7.52E+01 | 3.50E-45 | 0.058 | 2.16E+01 | 1.80E-19 | 0.020 |
| Limbic-FPN | 6.61E+02 | 0 | 0.350 | 3.07E+02 | 2.69E-287 | 0.223 |
| Limbic-DMN | 5.88E+02 | 2.13e-310 | 0.324 | 3.01E+02 | 2.64E-282 | 0.220 |
| FPN-DMN | 9.98E+02 | 0 | 0.448 | 6.09E+02 | 0 | 0.363 |

We performed one-way ANCOVAs of the factor connectome state on each FC_Time-Varying_ between ICN pairs, adjusted for age and sex. *K*: the chosen number of discrete connectome states, *P*^†^: *P* values Bonferroni-corrected for 105 tests (21 dependent variables of all ICN pairs and five frequency bands), *η^2^_p_*: Partial Eta squared effect size, n/a: not applicable.

### Table S2. ANCOVAs of the factor connectome state (*K* levels) on Modularity_Time-Varying_ and Fractional Occupancy

| **Variables** | ***K* = 4** |  |  | ***K* = 6** |  |  |
| --- | --- | --- | --- | --- | --- | --- |
|  | ***F* states** | ***P*^†^** | ***η^2^_p_*** | ***F* states** | ***P*^†^** | ***η^2^_p_*** |
| **Delta band** |  |  |  |  |  |  |
| Modularity_Time-Varying_ | 1.41E+02 | 2.68E-85 | 0.103 | 3.47E+02 | 0 | 0.238 |
| Fractional Occupancy | 5.16E+03 | 0 | 0.807 | 2.43E+03 | 0 | 0.686 |
| **Theta band** |  |  |  |  |  |  |
| Modularity_Time-Varying_ | 8.67E+01 | 6.33E-53 | 0.066 | 1.98E+02 | 3.16E-193 | 0.152 |
| Fractional Occupancy | 9.30E+02 | 0 | 0.430 | 1.16E+03 | 0 | 0.510 |
| **Alpha band** |  |  |  |  |  |  |
| Modularity_Time-Varying_ | 6.22E+01 | 6.38E-38 | 0.048 | 8.00E+01 | 5.11E-80 | 0.067 |
| Fractional Occupancy | 1.86E+03 | 0 | 0.601 | 1.14E+03 | 0 | 0.507 |
| **Beta band** |  |  |  |  |  |  |
| Modularity_Time-Varying_ | 3.30E+02 | 1.60E-188 | 0.212 | 1.08E+02 | 9.47E-107 | 0.105 |
| Fractional Occupancy | 7.20E+02 | 0 | 0.369 | 2.05E+03 | 0 | 0.649 |
| **Gamma band** |  |  |  |  |  |  |
| Modularity_Time-Varying_ | 1.55E+02 | 2.29E-93 | 0.112 | 6.48E+01 | 1.33E-64 | 0.057 |
| Fractional Occupancy | 5.44E+02 | 1.06E-29 | 0.306 | 8.89E+02 | 0 | 0.445 |

We performed one-way ANCOVAs of the factor connectome state on Modularity_Time-Varying_ and Fractional Occupancy, adjusted for age and sex. *K*: the chosen number of discrete connectome states, *P*^†^: *P* values Bonferroni-corrected for 20 tests (four main dependent variables and five frequency bands), *η^2^_p_*: Partial Eta squared effect size.

### Table S3. ANCOVAs of the factor sibling status (*3* levels) on Modularity_Time-Varying_ and FC_Time-Varying_ of data-driven clusters with different initial connection-wise thresholds (density).

|  | ***K = 6*** |  |  |  | ***K = 4*** | | | |
| --- | --- | --- | --- | --- | --- | --- | --- | --- |
| **Phenotypes** | ***F* sibling status** | ***P*^†^** | ***η^2^_p_*** | **BF_01_** | ***F* sibling status** | ***P*^†^** | ***η^2^_p_*** | **BF_01_** |
| **Delta (1-3 Hz)** |  |  |  |  |  |  |  |  |
| Modularity_Time-Varying_ | .08 | 1.00 | <.001 | 36.79 | 5.27 | .108 | .023 | .231 |
| NBS-FC_Time-Varying_ 5% | 1.122 | 1.00 | 0.005 | 13.81 | 0.869 | 1.00 | 0.004 | 17.65 |
| NBS-FC_Time-Varying_ 4% | n/a | n/a | n/a | n/a | 0.792 | 1.00 | 0.003 | 18.69 |
| NBS-FC_Time-Varying_ 3% | 1.981 | 1.00 | 0.009 | 5.72 | 0.72 | 1.00 | 0.003 | 20.08 |
| NBS-FC_Time-Varying_ 2% | 1.989 | 1.00 | 0.009 | 6.34 | 1.586 | 1.00 | 0.007 | 8.93 |
| NBS-FC_Time-Varying_ 1% | 1.152 | 1.00 | 0.005 | 13.79 | 0.004 | 1.00 | 1.67e-05 | 39.54 |
| **Theta (4-7 Hz)** |  |  |  |  |  |  |  |  |
| Modularity_Time-Varying_ | 2.88 | 1.00 | .012 | 2.90 | .768 | 1.00 | .003 | 17.79 |
| NBS-FC_Time-Varying_ 5% | 1.252 | 1.00 | 0.005 | 12.41 | 0.647 | 1.00 | 0.003 | 21.74 |
| NBS-FC_Time-Varying_ 4% | 1.529 | 1.00 | 0.007 | 9.61 | 0.393 | 1.00 | 0.002 | 27.42 |
| NBS-FC_Time-Varying_ 3% | 1.066 | 1.00 | 0.005 | 14.62 | 0.289 | 1.00 | 0.001 | 29.69 |
| NBS-FC_Time-Varying_ 2% | 0.719 | 1.00 | 0.003 | 19.90 | 0.86 | 1.00 | 0.004 | 17.45 |
| NBS-FC_Time-Varying_ 1% | 1.379 | 1.00 | 0.006 | 10.55 | 0.902 | 1.00 | 0.004 | 16.60 |
| **Alpha (8-12 Hz)** |  |  |  |  |  |  |  |  |
| Modularity_Time-Varying_ | 3.57 | .577 | .015 | 1.72 | .300 | 1.00 | .001 | 31.65 |
| NBS-FC_Time-Varying_ 5% | 0.181 | 1.00 | 7.91e-04 | 33.57 | 0.206 | 1.00 | 8.98e-04 | 32.61 |
| NBS-FC_Time-Varying_ 4% | 0.119 | 1.00 | 5.18e-04 | 35.66 | 0.129 | 1.00 | 5.61e-04 | 34.99 |
| NBS-FC_Time-Varying_ 3% | 0.11 | 1.00 | 4.82e-04 | 36.58 | 0.164 | 1.00 | 7.14e-04 | 33.30 |
| NBS-FC_Time-Varying_ 2% | n/a | n/a | n/a | n/a | 0.02 | 1.00 | 8.70e-05 | 38.49 |
| NBS-FC_Time-Varying_ 1% | 0.24 | 1.00 | 0.001 | 32.93 | 0.96 | 1.00 | 0.004 | 14.22 |
| **Beta (13-25 Hz)** |  |  |  |  |  |  |  |  |
| Modularity_Time-Varying_ | .93 | 1.00 | .004 | 17.05 | 1.05 | 1.00 | .005 | 14.97 |
| NBS-FC_Time-Varying_ 5% | 0.076 | 1.00 | 3.33e-04 | 37.01 | 1.314 | 1.00 | 0.006 | 11.52 |
| NBS-FC_Time-Varying_ 4% | 0.092 | 1.00 | 4.02e-04 | 36.48 | 1.749 | 1.00 | 0.008 | 7.65 |
| NBS-FC_Time-Varying_ 3% | 0.086 | 1.00 | 3.76e-04 | 36.85 | n/a | n/a | n/a | n/a |
| NBS-FC_Time-Varying_ 2% | 0.144 | 1.00 | 6.28e-04 | 35.16 | 2.581 | 1.00 | 0.011 | 3.74 |
| NBS-FC_Time-Varying_ 1% | 0.128 | 1.00 | 5.60e-04 | 36.05 | 0.558 | 1.00 | 0.002 | 23.41 |
| **Gamma (30-45 Hz)** |  |  |  |  |  |  |  |  |
| Modularity_Time-Varying_ | .45 | 1.00 | .002 | 23.88 | .126 | 1.00 | .001 | 35.25 |
| NBS-FC_Time-Varying_ 5% | 0.182 | 1.00 | 7.95e-04 | 33.64 | n/a | n/a | n/a | n/a |
| NBS-FC_Time-Varying_ 4% | n/a | n/a | n/a | n/a | 1.1 | 1.00 | 0.005 | 13.33 |
| NBS-FC_Time-Varying_ 3% | 0.201 | 1.00 | 8.76e-04 | 33.72 | 1.104 | 1.00 | 0.005 | 13.19 |
| NBS-FC_Time-Varying_ 2% | 0.158 | 1.00 | 6.88e-04 | 35.32 | 1.101 | 1.00 | 0.005 | 13.10 |
| NBS-FC_Time-Varying_ 1% | 0.502 | 1.00 | 0.002 | 24.93 | 2.067 | 1.00 | 0.009 | 5.38 |

We performed Network-Based Statistics (NBS) to select connected sets of connections (i.e., clusters) in a data-driven manner (Zalesky et al., 2010). We started with the absolute functional connectivity (FC) matrices, where all connections were transformed into absolute values to focus on the strength of connections, regardless of their connectivity direction. NBS implements an ANCOVA of the factor state for every connection and quantifies the degree to which each connection’s connectivity strength differed across states, adjusted for age and head motion. The ensuing connection-wise *F* statistics was thresholded at arbitrary values, allowing 1 ~ 5% of connections to survive. The multivariate spatial features were constructed as 1 × *K* vector by averaging the absolute FC values of all connections of the cluster for each state. *F* values are reported for one-way ANCOVAs of the factor sibling status (three levels: monozygotic twins (MZ), sex-matched dizygotic twins (DZ), and sex-matched pairs of unrelated individuals), adjusted for age and sex. *K*: the chosen number of discrete connectome states, *P*^†^: *P* values Bonferroni-corrected for 25 tests for NBS-FC_Time-Varying_ measures (five density levels and five frequency bands; capped at 1 maximum) and for 20 tests for Modularity_Time-Varying_ (four main multivariate features and five frequency bands), *η^2^_p_*: Partial Eta squared effect size, BF_01_: the probability of H_0_ against H_1_.

### Table S4. ANCOVAs of the factor sibling status (*3* levels) on FC_Time-Varying_ of all Intrinsic connectivity network (ICN) pairs.

| **Variables** | ***K* = 4** | | | ***K* = 6** | | |
| --- | --- | --- | --- | --- | --- | --- |
|  | ***F* sibling status** | ***P*^†^** | ***η^2^_p_*** | ***F* sibling status** | ***P*^†^** | ***η^2^_p_*** |
| **Delta band** |  |  |  |  |  |  |
| VIS-SMN_Time-Varying_ | 0.259 | 1 | 0.001 | 0.358 | 1 | 0.002 |
| VIS-DAN_Time-Varying_ | 1.146 | 1 | 0.005 | 0.215 | 1 | 9.39e-04 |
| VIS-VAN_Time-Varying_ | 0.031 | 1 | 1.36e-04 | 0.148 | 1 | 6.47e-04 |
| VIS-Limbic_Time-Varying_ | 0.132 | 1 | 5.77e-04 | 0.323 | 1 | 0.001 |
| VIS-FPN_Time-Varying_ | 0.298 | 1 | 0.001 | 0.088 | 1 | 3.84e-04 |
| VIS-DMN_Time-Varying_ | 0.633 | 1 | 0.003 | 0.65 | 1 | 0.003 |
| SMN-DAN_Time-Varying_ | 1.213 | 1 | 0.005 | 0.632 | 1 | 0.003 |
| SMN-VAN_Time-Varying_ | 0.454 | 1 | 0.002 | 0.125 | 1 | 5.47e-04 |
| SMN-Limbic_Time-Varying_ | 2.168 | 1 | 0.009 | 0.428 | 1 | 0.002 |
| SMN-FPN_Time-Varying_ | 0.201 | 1 | 8.80e-04 | 1.133 | 1 | 0.005 |
| SMN-DMN_Time-Varying_ | 1.524 | 1 | 0.007 | 1.613 | 1 | 0.007 |
| DAN-VAN_Time-Varying_ | 0.327 | 1 | 0.001 | 1.325 | 1 | 0.006 |
| DAN-Limbic_Time-Varying_ | 0.372 | 1 | 0.002 | 0.262 | 1 | 0.001 |
| DAN-FPN_Time-Varying_ | 0.369 | 1 | 0.002 | 0.073 | 1 | 3.17e-04 |
| DAN-DMN_Time-Varying_ | 0.257 | 1 | 0.001 | 0.904 | 1 | 0.004 |
| VAN-Limbic_Time-Varying_ | 0.993 | 1 | 0.004 | 0.926 | 1 | 0.004 |
| VAN-FPN_Time-Varying_ | 0.475 | 1 | 0.002 | 0.86 | 1 | 0.004 |
| VAN-DMN_Time-Varying_ | 0.963 | 1 | 0.004 | 3.003 | 1 | 0.013 |
| Limbic-FPN_Time-Varying_ | 0.548 | 1 | 0.002 | 0.109 | 1 | 4.77e-04 |
| Limbic-DMN_Time-Varying_ | 0.636 | 1 | 0.003 | 0.566 | 1 | 0.002 |
| FPN-DMN_Time-Varying_ | 0.194 | 1 | 8.52e-04 | 0.194 | 1 | 8.45e-04 |
| **Theta band** |  |  |  |  |  |  |
| VIS-SMN_Time-Varying_ | 2.354 | 1 | 0.01 | 3.185 | 1 | 0.014 |
| VIS-DAN_Time-Varying_ | 0.029 | 1 | 1.29e-04 | 0.074 | 1 | 3.25e-04 |
| VIS-VAN_Time-Varying_ | 0.019 | 1 | 8.57e-05 | 1.089 | 1 | 0.005 |
| VIS-Limbic_Time-Varying_ | 1.088 | 1 | 0.005 | 0.674 | 1 | 0.003 |
| VIS-FPN_Time-Varying_ | 0.279 | 1 | 0.001 | 0.573 | 1 | 0.002 |
| VIS-DMN_Time-Varying_ | 1.416 | 1 | 0.006 | 1.909 | 1 | 0.008 |
| SMN-DAN_Time-Varying_ | 0.317 | 1 | 0.001 | 0.361 | 1 | 0.002 |
| SMN-VAN_Time-Varying_ | 0.255 | 1 | 0.001 | 0.986 | 1 | 0.004 |
| SMN-Limbic_Time-Varying_ | 2.478 | 1 | 0.011 | 0.71 | 1 | 0.003 |
| SMN-FPN_Time-Varying_ | 0.147 | 1 | 6.52e-04 | 0.069 | 1 | 3.01e-04 |
| SMN-DMN_Time-Varying_ | 0.174 | 1 | 7.70e-04 | 0.175 | 1 | 7.62e-04 |
| DAN-VAN_Time-Varying_ | 1.339 | 1 | 0.006 | 0.812 | 1 | 0.004 |
| DAN-Limbic_Time-Varying_ | 1.864 | 1 | 0.008 | 1.878 | 1 | 0.008 |
| DAN-FPN_Time-Varying_ | 0.225 | 1 | 9.94e-04 | 0.413 | 1 | 0.002 |
| DAN-DMN_Time-Varying_ | 0.825 | 1 | 0.004 | 0.241 | 1 | 0.001 |
| VAN-Limbic_Time-Varying_ | 0.631 | 1 | 0.003 | 0.096 | 1 | 4.20e-04 |
| VAN-FPN_Time-Varying_ | 0.482 | 1 | 0.002 | 0.446 | 1 | 0.002 |
| VAN-DMN_Time-Varying_ | 0.2 | 1 | 8.84e-04 | 0.488 | 1 | 0.002 |
| Limbic-FPN_Time-Varying_ | 2.413 | 1 | 0.011 | 0.051 | 1 | 2.23e-04 |
| Limbic-DMN | 0.072 | 1 | 3.18e-04 | 1.063 | 1 | 0.005 |
| FPN-DMN | 0.149 | 1 | 6.57e-04 | 0.706 | 1 | 0.003 |
| **Alpha band** |  |  |  |  |  |  |
| VIS-SMN | 3.577 | 1 | 0.016 | 0.413 | 1 | 0.002 |
| VIS-DAN | 0.172 | 1 | 7.60e-04 | 0.454 | 1 | 0.002 |
| VIS-VAN | 6.649 | 0 | 0.029 | 0.172 | 1 | 7.51e-04 |
| VIS-Limbic | 4.54 | 1 | 0.02 | 0.041 | 1 | 1.79e-04 |
| VIS-FPN | 3.29 | 1 | 0.014 | 0.453 | 1 | 0.002 |
| VIS-DMN | 5.158 | 0.6 | 0.022 | 1.278 | 1 | 0.006 |
| SMN-DAN | 1.451 | 1 | 0.006 | 0.851 | 1 | 0.004 |
| SMN-VAN | 1.886 | 1 | 0.008 | 1.379 | 1 | 0.006 |
| SMN-Limbic | 2.299 | 1 | 0.01 | 0.475 | 1 | 0.002 |
| SMN-FPN | 3.041 | 1 | 0.013 | 1.106 | 1 | 0.005 |
| SMN-DMN | 4.311 | 1 | 0.019 | 3.599 | 1 | 0.015 |
| DAN-VAN | 0.463 | 1 | 0.002 | 0.738 | 1 | 0.003 |
| DAN-Limbic | 0.046 | 1 | 2.03e-04 | 1.068 | 1 | 0.005 |
| DAN-FPN | 0.907 | 1 | 0.004 | 1.015 | 1 | 0.004 |
| DAN-DMN | 1.799 | 1 | 0.008 | 0.12 | 1 | 5.25e-04 |
| VAN-Limbic | 0.514 | 1 | 0.002 | 0.72 | 1 | 0.003 |
| VAN-FPN | 1.033 | 1 | 0.005 | 0.74 | 1 | 0.003 |
| VAN-DMN | 5.086 | 0.6 | 0.022 | 1.741 | 1 | 0.008 |
| Limbic-FPN | 1.854 | 1 | 0.008 | 0.724 | 1 | 0.003 |
| Limbic-DMN | 4.73 | 1 | 0.02 | 0.018 | 1 | 7.94e-05 |
| FPN-DMN | 1.214 | 1 | 0.005 | 0.185 | 1 | 8.06e-04 |
| **Beta band** |  |  |  |  |  |  |
| VIS-SMN | 2.016 | 1 | 0.009 | 1.933 | 1 | 0.008 |
| VIS-DAN | 1.149 | 1 | 0.005 | 0.344 | 1 | 0.001 |
| VIS-VAN | 1.104 | 1 | 0.005 | 0.209 | 1 | 9.11e-04 |
| VIS-Limbic | 1.613 | 1 | 0.007 | 0.54 | 1 | 0.002 |
| VIS-FPN | 1.199 | 1 | 0.005 | 0.605 | 1 | 0.003 |
| VIS-DMN | 1.792 | 1 | 0.008 | 1.378 | 1 | 0.006 |
| SMN-DAN | 1.987 | 1 | 0.009 | n/a | n/a | n/a |
| SMN-VAN | 1.072 | 1 | 0.005 | 0.053 | 1 | 2.29e-04 |
| SMN-Limbic | 0.319 | 1 | 0.001 | 0.056 | 1 | 2.43e-04 |
| SMN-FPN | 0.378 | 1 | 0.002 | n/a | n/a | n/a |
| SMN-DMN | 0.312 | 1 | 0.001 | 0.878 | 1 | 0.004 |
| DAN-VAN | 1.628 | 1 | 0.007 | n/a | n/a | n/a |
| DAN-Limbic | 0.04 | 1 | 1.78e-04 | n/a | n/a | n/a |
| DAN-FPN | 1.474 | 1 | 0.006 | n/a | n/a | n/a |
| DAN-DMN | 0.036 | 1 | 1.60e-04 | 0.37 | 1 | 0.002 |
| VAN-Limbic | 1.39 | 1 | 0.006 | 0.119 | 1 | 5.17e-04 |
| VAN-FPN | 0.453 | 1 | 0.002 | 1.46 | 1 | 0.006 |
| VAN-DMN | 0.346 | 1 | 0.002 | 1.014 | 1 | 0.004 |
| Limbic-FPN | 0.19 | 1 | 8.42e-04 | 2.052 | 1 | 0.009 |
| Limbic-DMN | 0.231 | 1 | 0.001 | 0.183 | 1 | 7.99e-04 |
| FPN-DMN | 0.03 | 1 | 1.31e-04 | 0.522 | 1 | 0.002 |
| **Gamma band** |  |  |  |  |  |  |
| VIS-SMN | 2.118 | 1 | 0.009 | 0.383 | 1 | 0.002 |
| VIS-DAN | 1.381 | 1 | 0.006 | 0.125 | 1 | 5.46e-04 |
| VIS-VAN | 0.516 | 1 | 0.002 | 5.537 | 0.6 | 0.024 |
| VIS-Limbic | 0.909 | 1 | 0.004 | 1.148 | 1 | 0.005 |
| VIS-FPN | 3.499 | 1 | 0.015 | 1.027 | 1 | 0.004 |
| VIS-DMN | 1.591 | 1 | 0.007 | 0.956 | 1 | 0.004 |
| SMN-DAN | 0.883 | 1 | 0.004 | 0.091 | 1 | 3.96e-04 |
| SMN-VAN | 0.134 | 1 | 5.85e-04 | 0.303 | 1 | 0.001 |
| SMN-Limbic | 1.044 | 1 | 0.005 | 0.154 | 1 | 6.71e-04 |
| SMN-FPN | 0.862 | 1 | 0.004 | 0.401 | 1 | 0.002 |
| SMN-DMN | 1.189 | 1 | 0.005 | 0.475 | 1 | 0.002 |
| DAN-VAN | 2.565 | 1 | 0.011 | 2.174 | 1 | 0.009 |
| DAN-Limbic | 0.692 | 1 | 0.003 | 0.383 | 1 | 0.002 |
| DAN-FPN | 2.293 | 1 | 0.01 | n/a | n/a | n/a |
| DAN-DMN | 2.021 | 1 | 0.009 | 1.438 | 1 | 0.006 |
| VAN-Limbic | 1.438 | 1 | 0.006 | 0.695 | 1 | 0.003 |
| VAN-FPN | 1.925 | 1 | 0.008 | 1.055 | 1 | 0.005 |
| VAN-DMN | 0.955 | 1 | 0.004 | 2.546 | 1 | 0.011 |
| Limbic-FPN | 0.854 | 1 | 0.004 | 0.078 | 1 | 3.39e-04 |
| Limbic-DMN | 0.234 | 1 | 0.001 | 2.408 | 1 | 0.01 |
| FPN-DMN | 4.815 | 1 | 0.021 | 0.944 | 1 | 0.004 |

Heritability of each of the FC_Time-Varying_ between ICN pairs was assessed separately by one-way ANCOVAs of the factor sibling status (three levels: monozygotic twins (MZ), sex-matched dizygotic twins (DZ), and sex-matched pairs of unrelated individuals), adjusted for age and sex. The main effect of sibling status indicates the heritability, or genetic effect. *K*: the chosen number of discrete connectome states, *P*^†^: *P* values Bonferroni-corrected for 105 tests (5 frequency bands and 21 dependent variables of all ICN pairs), *η^2^_p_*: Partial Eta squared effect size.

### Table S5. Heritability of individual components of temporal dynamic connectome features.

| **Frequency band** | **Variables** | ***F* sibling status** | ***P*^†^ value** | ***η^2^_p_*** | ***F interaction*** | ***P*^†^ value** |
| --- | --- | --- | --- | --- | --- | --- |
| ***K* = 4** |  |  |  |  |  |  |
| Delta band | Transition Probability | 11.84 | 7.39e-05 | 0.004 | 1.83 | .104 |
|  | Fractional Occupancy | 12.06 | 6.29e-05 | 0.013 | .207 | 1.00 |
| Theta band | Transition Probability | 21.79 | 3.76e-09 | 0.008 | 2.66 | 4.07e-04 |
|  | Fractional Occupancy | 19.73 | 3.34e-08 | 0.021 | .691 | 1.00 |
| Alpha band | Transition Probability | 121.08 | 3.50e-51 | 0.042 | 2.87 | 8.44e-05 |
|  | Fractional Occupancy | 23.29 | 1.03e-09 | 0.025 | .951 | 1.00 |
| Beta band | Transition Probability | 38.37 | 2.83e-16 | 0.014 | 4.09 | 4.24e-09 |
|  | Fractional Occupancy | 24.18 | 4.33e-10 | 0.026 | 2.02 | .603 |
| Gamma band | Transition Probability | 27.68 | 1.09e-11 | 0.01 | 4.09 | 4.31e-09 |
|  | Fractional Occupancy | 23.45 | 8.81e-10 | 0.025 | 2.77 | .111 |
| ***K* = 6** |  |  |  |  |  |  |
| Delta band | Transition Probability | 23.17 | 9.03e-10 | 0.003 | 1.39 | .264 |
|  | Fractional Occupancy | 17.32 | 3.34e-07 | 0.012 | .621 | 1 |
| Theta band | Transition Probability | 89.08 | 3.67e-38 | 0.013 | 1.37 | .314 |
|  | Fractional Occupancy | 33.34 | 4.92e-14 | 0.024 | 1.23 | 1 |
| Alpha band | Transition Probability | 127.01 | 2.20e-54 | 0.018 | 3.61 | 8.53e-18 |
|  | Fractional Occupancy | 51.79 | 8.33e-22 | 0.036 | 2.29 | .114 |
| Beta band | Transition Probability | 32.61 | 7.44e-14 | 0.005 | 3.81 | 1.28e-19 |
|  | Fractional Occupancy | 25.37 | 1.21e-10 | 0.018 | 3.60 | 9.16e-04 |
| Gamma band | Transition Probability | 69.27 | 1.16e-29 | 0.01 | 2.52 | 1.77e-08 |
|  | Fractional Occupancy | 25.19 | 1.44e-10 | 0.018 | 3.44 | 1.69e-03 |

A two-way ANCOVA was conducted to examine the effect of sibling status on the *state-wise* similarity between subject pairs for each temporal feature. Specifically, the factors comprised sibling status (four levels: monozygotic twins (MZ), sex-matched dizygotic twins (DZ), and sex-matched pairs of unrelated individuals) and connectome states or state pairs (*K=*4 or 6 states for Fractional Occupancy and 12 or 30 state pairs for Transition Probability, respectively), adjusted for age and sex. *F* values are reported for the factor sibling status. The effect sizes (η^2^_p_) for the factor sibling status were smaller than those of the multivariate temporal features in the main analyses (cf. Figure 3). Interestingly, Transition Probability showed significant interaction effects, irrespective of the number of Ks and most of the frequency bands. This implies that heritability is dependent on specific pair of Transition Probability between connectome states. For completeness, we also note that, non-surprisingly, there were significant main effects of the factor connectome states, implying that the similarity across pairs of subjects varied by state. K: the chosen number of discrete connectome states, *P*^†^: *P* values Bonferroni-corrected for 10 tests (2 dependent variables and five frequency bands), *η^2^_p_*: Partial Eta squared effect size of the sibling status.

### Table S6. Variance-component model parameter estimates of the dynamic connectome features (K = 4).

| **Phenotypes** | **Genetic**  **model** | ***h^2^*** | **(95% CI)** | **A** | **(95% CI)** | **C/D** | **(95% CI)** | **E** | **(95% CI)** | **-2LL** | **AIC** | **chi** | **df** | **p(chi)** |
| --- | --- | --- | --- | --- | --- | --- | --- | --- | --- | --- | --- | --- | --- | --- |
| **Delta band** |  |  |  |  |  |  |  |  |  |  |  |  |  |  |
| Fractional Occupancy | **ADE** | **0.157** | **(.03, .29)** | 0.00 | (.00, .00) | 0.16 | (.03, .29) | 0.84 | (.71, .97) | -2563.9 | -2551.9 |  |  |  |
|  | AE | 0.148 | (.02, .27) | 0.15 | (.02, .27) |  |  | 0.85 | (.73, .98) | -2563.7 | -2553.7 | 0.2 | 1 | 0.664 |
|  | DE | 0.157 | (.03, .29) |  |  | 0.16 | (.03, .29) | 0.84 | (.71, .97) | -2563.9 | -2553.9 | 0.2 | 1 | 0.664 |
| Transition Probability | ACE | 0 | (.00, .00) | 0.00 | (.00, .00) | 0.09 | (.02, .20) | 0.91 | (.80, 1.02) | -2024.1 | -2012.1 |  |  |  |
|  | AE | 0.088 | (-.03, .21) | 0.09 | (.03, .21) |  |  | 0.91 | (.79, 1.03) | -2023.3 | -2013.3 | 0.7 | 1 | 0.387 |
|  | CE | 0 | (.00, .00) |  |  | 0.09 | (.02, .20) | 0.91 | (.80, 1.02) | -2024.1 | -2014.1 | 0.7 | 1 | 0.387 |
| Modularity_Time-Varying_ | ACE | 0 | (.00, .00) | 0.00 | (.00, .00) | 0.05 | (.06, .16) | 0.95 | (.84, 1.06) | -14.6 | -128.6 |  |  |  |
|  | AE | 0.034 | (-.10, .16) | 0.03 | (.10, .16) |  |  | 0.97 | (.84, 1.10) | -14.2 | -13.2 | 0.4 | 1 | 0.523 |
|  | CE | 0 | (.00, .00) |  |  | 0.05 | (.06, .16) | 0.95 | (.84, 1.06) | -14.6 | -13.6 | 0.4 | 1 | 0.523 |
| FC_Time-Varying_ of  data-driven clusters (4%) | ADE | 0.035 | (-.05, .11) | 0.00 | (.00, .00) | 0.04 | (.05, .11) | 0.97 | (.89, 1.05) | -7003.4 | -6991.4 |  |  |  |
|  | AE | 0.031 | (-.05, .11) | 0.03 | (.05, .11) |  |  | 0.97 | (.89, 1.05) | -7003.4 | -6993.4 | 0.0 | 1 | 0.838 |
|  | DE | 0.035 | (-.05, .11) |  |  | 0.04 | (.05, .11) | 0.97 | (.89, 1.05) | -7003.4 | -6993.4 | 0.0 | 1 | 0.838 |
| **Theta band** |  |  |  |  |  |  |  |  |  |  |  |  |  |  |
| Fractional Occupancy | ACE | 0.166 | (-.21, .54) | 0.17 | (.21, .54) | 0.05 | (.27, .37) | 0.79 | (.66, .91) | -2493.0 | -249.0 |  |  |  |
|  | AE | 0.219 | (.10, .34) | 0.22 | (.10, .34) |  |  | 0.78 | (.66, .90) | -2492.9 | -2482.9 | 0.1 | 1 | 0.804 |
|  | CE | 0 | (.00, .00) |  |  | 0.18 | (.08, .29) | 0.82 | (.71, .92) | -2492.4 | -2482.4 | 0.1 | 1 | 0.804 |
| Transition Probability | ACE | 0.071 | (-.33, .47) | 0.07 | (.33, .47) | 0.04 | (.29, .38) | 0.89 | (.75, 1.02) | -1584.1 | -1572.1 |  |  |  |
|  | AE | 0.121 | (-.01, .25) | 0.12 | (.01, .25) |  |  | 0.88 | (.75, 1.01) | -1584.0 | -1574.0 | 0.1 | 1 | 0.82 |
|  | CE | 0 | (.00, .00) |  |  | 0.10 | (.01, .21) | 0.90 | (.79, 1.01) | -1584.0 | -1574.0 | 0.1 | 1 | 0.82 |
| Modularity_Time-Varying_ | ACE | 0 | (.00, .00) | 0.00 | (.00, .00) | 0.02 | (.09, .13) | 0.98 | (.87, 1.09) | 145.6 | 157.6 |  |  |  |
|  | AE | 0.003 | (-.12, .12) | 0.00 | (.12, .12) |  |  | 1.00 | (.88, 1.12) | 145.7 | 155.7 | 0.1 | 1 | 0.78 |
|  | CE | 0 | (.00, .00) |  |  | 0.02 | (.09, .13) | 0.98 | (.87, 1.09) | 145.6 | 155.6 | 0.1 | 1 | 0.78 |
| FC_Time-Varying_ of  data-driven clusters (4%) | ACE | 0 | (-.00, .00) | 0.00 | (.00, .00) | 0.05 | (.04, .13) | 0.96 | (.87, 1.04) | -6897.2 | -6885.2 |  |  |  |
|  | AE | 0.039 | (-.04, .12) | 0.04 | (.04, .12) |  |  | 0.96 | (.88, 1.04) | -6897.0 | -6887.0 | 0.2 | 1 | 0.685 |
|  | CE | 0 | (.00, .00) |  |  | 0.05 | (.03, .13) | 0.96 | (.87, 1.03) | -6897.2 | -6887.2 | 0.2 | 1 | 0.685 |
| **Alpha band** |  |  |  |  |  |  |  |  |  |  |  |  |  |  |
| Fractional Occupancy | ACE | 0 | (.00, .00) | 0.00 | (.00, .00) | 0.00 | (.00, .00) | 1.00 | (1.00, 1.00) | -1903.0 | -1891.0 |  |  |  |
|  | AE | 0 | (.00, .00) | 0.00 | (.00, .00) |  |  | 1.00 | (1.00, 1.00) | -1903.0 | -1893.0 | 6.8 | 1 | 1 |
|  | CE | 0 | (.00, .00) |  |  | 0.00 | (.00, .00) | 1.00 | (1.00, 1.00) | -1903.0 | -1893.0 | 6.8 | 1 | 1 |
| Transition Probability | ACE | 0.347 | (-.01, .71) | 0.35 | (.01, .71) | 0.18 | (.15, .51) | 0.47 | (.38, .57) | -1969.3 | -1957.3 |  |  |  |
|  | AE | 0.534 | (.45, .62) | 0.53 | (.45, .62) |  |  | 0.47 | (.38, .55) | -1968.4 | -1958.4 | 0.9 | 1 | 0.331 |
|  | CE | 0 | (.00, .00) |  |  | 0.47 | (.39, .56) | 0.53 | (.44, .61) | -1965.4 | -1955.4 | 0.9 | 1 | 0.331 |
| Modularity_Time-Varying_ | ADE | 0.129 | (-.00, .26) | 0.00 | (.00, .00) | 0.13 | (.00, .26) | 0.87 | (.74, 1.00) | 672.9 | 684.9 |  |  |  |
|  | AE | 0.12 | (-.01, .25) | 0.12 | (.01, .25) |  |  | 0.88 | (.75, 1.01) | 673.1 | 683.1 | 0.2 | 1 | 0.654 |
|  | DE | 0.129 | (-.00, .26) |  |  | 0.13 | (.00, .26) | 0.87 | (.74, 1.00) | 672.9 | 682.9 | 0.2 | 1 | 0.654 |
| FC_Time-Varying_ of  data-driven clusters (4%) | ADE | 0.075 | (-.03, .18) | 0.00 | (.00, .00) | 0.08 | (.03, .18) | 0.93 | (.82, 1.03) | -6546.7 | -6534.7 |  |  |  |
|  | AE | 0.064 | (-.03, .16) | 0.06 | (.03, .16) |  |  | 0.94 | (.84, 1.03) | -6546.3 | -6536.3 | 0.3 | 1 | 0.562 |
|  | DE | 0.075 | (-.02, .17) |  |  | 0.08 | (.02, .17) | 0.93 | (.83, 1.02) | -6546.7 | -6536.7 | 0.3 | 1 | 0.562 |
| **Beta band** |  |  |  |  |  |  |  |  |  |  |  |  |  |  |
| Fractional Occupancy | **ADE** | **0.307** | **(.17, .44)** | 0.10 | (.43, .62) | 0.21 | (.35, .77) | 0.69 | (.56, .83) | -2238.7 | -2226.7 |  |  |  |
|  | AE | 0.289 | (.16, .42) | 0.29 | (.16, .42) |  |  | 0.71 | (.58, .84) | -2238.3 | -2228.3 | 0.4 | 1 | 0.539 |
|  | DE | 0.311 | (.18, .44) |  |  | 0.31 | (.18, .44) | 0.69 | (.56, .82) | -2238.6 | -2228.6 | 0.4 | 1 | 0.539 |
| Transition Probability | **ADE** | **0.198** | **(.05, .34)** | 0.00 | (.00, .00) | 0.20 | (.05, .34) | 0.80 | (.66, .95) | -2204.5 | -2192.5 |  |  |  |
|  | AE | 0.166 | (.03, .30) | 0.17 | (.03, .30) |  |  | 0.83 | (.70, .97) | -2203.3 | -2193.3 | 1.2 | 1 | 0.268 |
|  | DE | 0.198 | (.05, .34) |  |  | 0.20 | (.05, .34) | 0.80 | (.66, .95) | -2204.5 | -2194.5 | 1.2 | 1 | 0.268 |
| Modularity_Time-Varying_ | ACE | 0.017 | (-.42, .45) | 0.02 | (.42, .45) | 0.09 | (.29, .46) | 0.89 | (.76, 1.03) | 515.9 | 527.9 |  |  |  |
|  | AE | 0.117 | (-.01, .24) | 0.12 | (.01, .24) |  |  | 0.88 | (.76, 1.01) | 516.1 | 526.1 | 0.2 | 1 | 0.654 |
|  | CE | 0 | (.00, .00) |  |  | 0.10 | (.01, .21) | 0.90 | (.79, 1.01) | 515.9 | 525.9 | 0.2 | 1 | 0.654 |
| FC_Time-Varying_ of  data-driven clusters (4%) | **ADE** | **0.124** | **(.01, .24)** | 0.00 | (.00, .00) | 0.12 | (.01, .24) | 0.88 | (.76, .99) | -6879.2 | -6867.2 |  |  |  |
|  | AE | 0.105 | (-.00, .21) | 0.11 | (.00, .21) |  |  | 0.90 | (.79, 1.00) | -6878.5 | -6868.5 | 0.7 | 1 | 0.394 |
|  | DE | 0.124 | (.01, .23) |  |  | 0.12 | (.01, .23) | 0.88 | (.77, .99) | -6879.2 | -6869.2 | 0.7 | 1 | 0.394 |
| **Gamma band** |  |  |  |  |  |  |  |  |  |  |  |  |  |  |
| Fractional Occupancy | **ADE** | **0.205** | **(.06, .35)** | 0.00 | (.00, .00) | 0.21 | (.06, .35) | 0.80 | (.65, .94) | -1886.5 | -1874.5 |  |  |  |
|  | AE | 0.174 | (.03, .31) | 0.17 | (.03, .31) |  |  | 0.83 | (.69, .97) | -1885.6 | -1875.6 | 0.9 | 1 | 0.33 |
|  | DE | 0.205 | (.06, .35) |  |  | 0.21 | (.06, .35) | 0.80 | (.65, .94) | -1886.5 | -1876.5 | 0.9 | 1 | 0.33 |
| Transition Probability | **ADE** | **0.318** | **(.20, .44)** | 0.00 | (.00, .00) | 0.32 | (.20, .44) | 0.68 | (.56, .80) | -1581.9 | -1569.9 |  |  |  |
|  | AE | 0.298 | (.18, .41) | 0.30 | (.18, .41) |  |  | 0.70 | (.59, .82) | -158.2 | -157.2 | 1.6 | 1 | 0.202 |
|  | DE | 0.318 | (.20, .44) |  |  | 0.32 | (.20, .44) | 0.68 | (.56, .80) | -1581.9 | -1571.9 | 1.6 | 1 | 0.202 |
| Modularity_Time-Varying_ | ADE | 0.012 | (-.13, .15) | 0.00 | (.00, .00) | 0.01 | (.13, .15) | 0.99 | (.85, 1.13) | 659.0 | 671.0 |  |  |  |
|  | AE | 0 | (.00, .00) | 0.00 | (.00, .00) |  |  | 1.00 | (1.00, 1.00) | 659.1 | 669.1 | 0.0 | 1 | 0.86 |
|  | DE | 0.012 | (-.13, .15) |  |  | 0.01 | (.13, .15) | 0.99 | (.85, 1.13) | 659.0 | 669.0 | 0.0 | 1 | 0.86 |
| FC_Time-Varying_ of  data-driven clusters (4%) | ACE | 0 | (.00, .00) | 0.00 | (.00, .00) | 0.00 | (.00, .00) | 1.00 | (1.00, 1.00) | -6539.5 | -6527.5 |  |  |  |
|  | AE | 0 | (.00, .00) | 1.00 | (1.00, 1.00) |  |  | 1.00 | (1.00, 1.00) | -6539.5 | -6529.5 | 7.9 | 9 | 1 |
|  | CE | 0 | (.00, .00) |  |  | 0.00 | (.00, .00) | 1.00 | (1.00, 1.00) | -6539.5 | -6529.5 | 7.9 | 9 | 1 |

All models were adjusted for age and sex. A, Additive genetic effect; C, common environmental effect; D, dominant genetic effect; E, Unique/non-shared environment effect; -2LL, twice the negative log-likelihood; AIC, Akaike’s information criterion; df, degrees of freedom; χ^2^, chi square, ∆df, change in degree of freedom between the full model and the nested model; *p*, χ^2^ test in model fitting. The AE and CE models are nested within the ACE model. Each nested model is compared with the fully saturated model. The fitness of models was tested based on a change in AIC (for a change of df of 1, the statistically significant change in χ^2^ is 3.84). *h^2^*, the narrow-sense heritability estimated as σ^2^_A_/(σ^2^_A_ + σ^2^_c_ + σ^2^_E_); the broad-sense heritability estimated as (σ^2^_A_ + σ^2^_D_)/(σ^2^_A_ + σ^2^_D_ + σ^2^_E_); CI: Confidence Interval (lower bound, upper bound).

**Supplementary Figures**

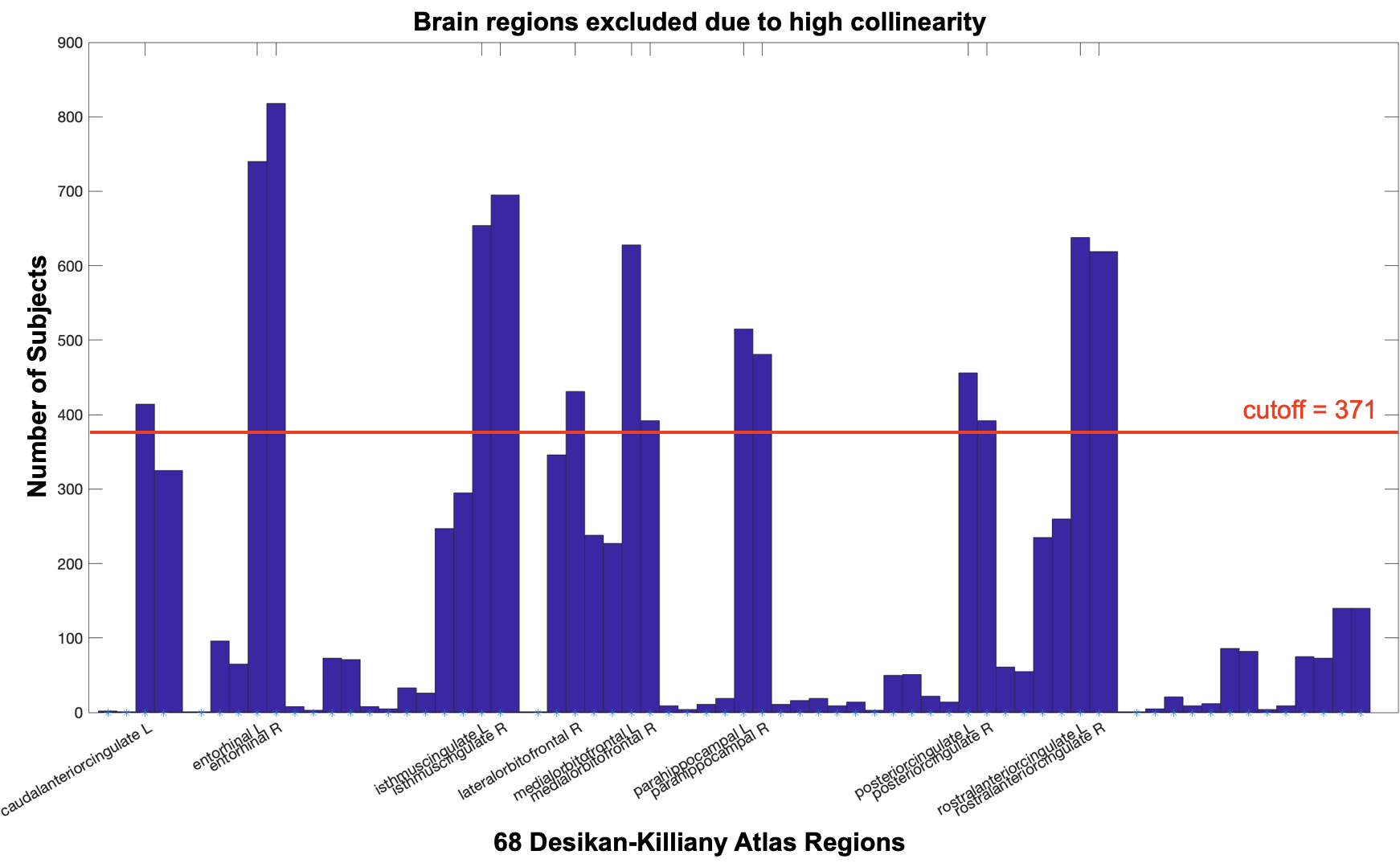

##### Figure S1. Top 14 regions with high collinearity.

To mitigate the source-leakage confound, caused by the blurring of point dipole sources and the spreading of signals across neighboring regions, we excluded regions, whose extracted signals were found to be highly collinear with others based on *qr* function in Matlab. Subsequently, we excluded 14 regions which displayed highly collinearity in over 40% of the 928 subjects (cutoff = 371) from the investigation: bilateral ‘*rostralanteriorcingulate’*, bilateral '*posteriorcingulate*', bilateral '*parahippocampal*', bilateral '*medialorbitofrontal'*, bilateral '*isthmuscingulate*', bilateral '*entorhinal*', '*lateralorbitofrontal R*', and '*caudalanteriorcingulate L'*.

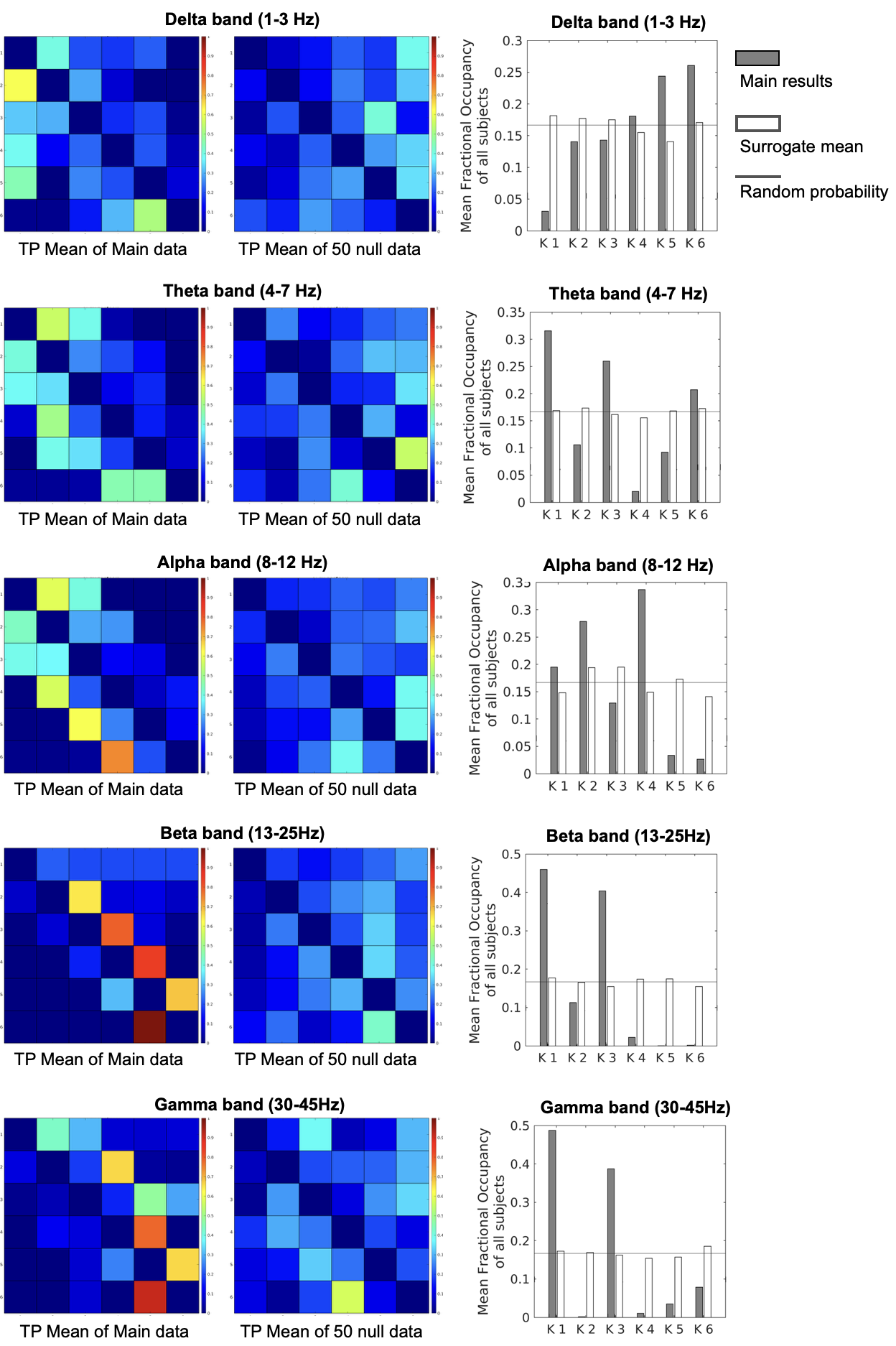

##### Figure S2. Difference in Transition Probability and Fractional Occupancy distribution obtained at *K* of 6 between main dataset and 50 surrogate datasets.

Fractional Occupancy and Transition Probability matrix averaged from the 50 surrogate data show little structured distribution across states (or state pairs), indicating the absence of meaningful dynamic properties in the surrogate data, whereas Fractional Occupancy and Transition Probability obtained from the real data show non-random sequencing of brain states.

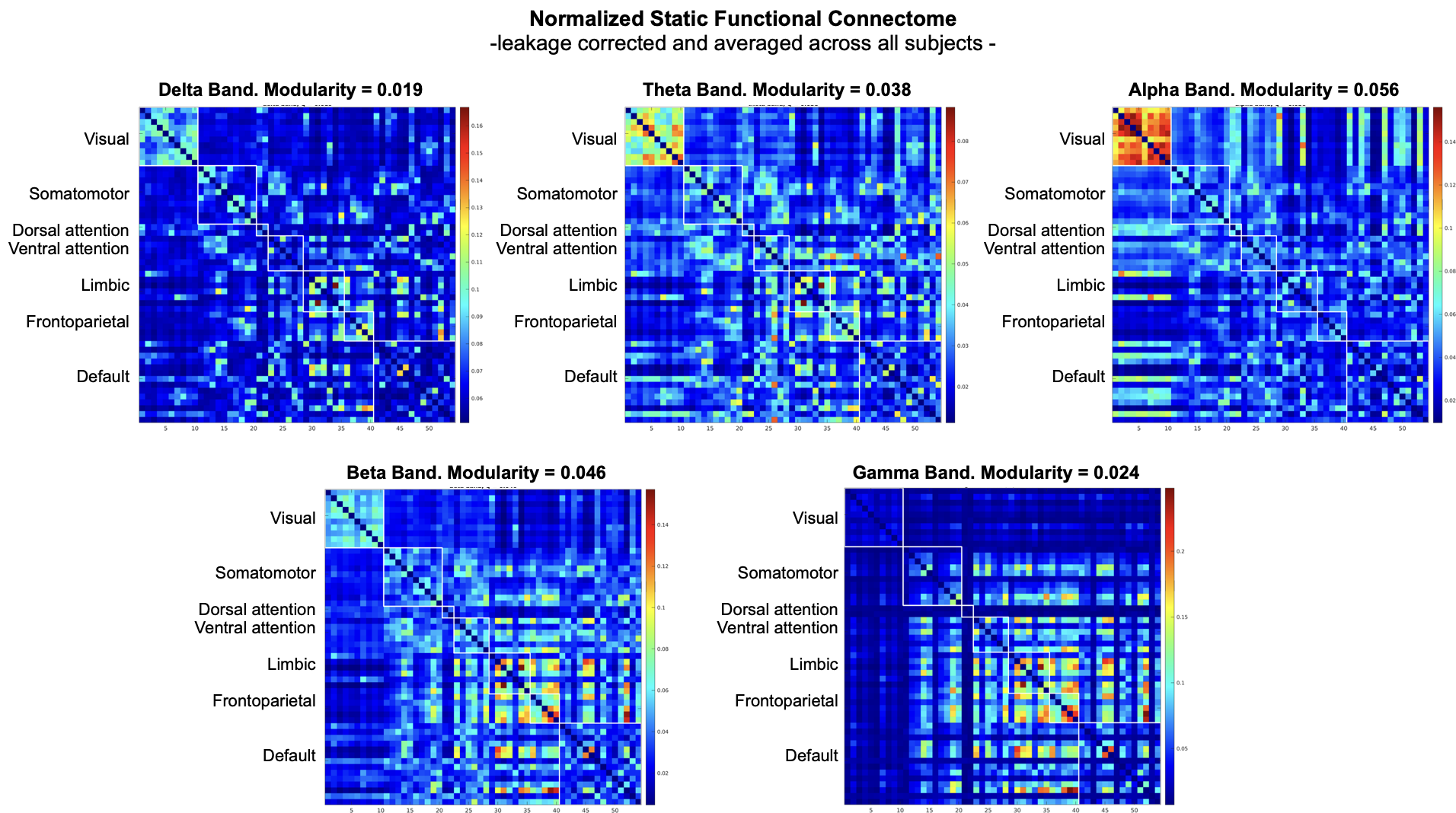

##### Figure S3. Static functional connectome for each canonical band averaged across all subjects in the study.

The 54 distinct regions of Desikan-Killiany Atlas, excluding 14 regions with high collinearity. The regional signals underwent detrending and bandpass filtering within canonical frequency ranges: delta (1-3 Hz), theta (4-7 Hz), alpha (8-12 Hz), beta (13-25 Hz), and gamma (30-45 Hz). Subsequently, we used a symmetric orthogonalization procedure (Colclough et al., 2015) to remove all shared signal at zero lag between the regions. Functional connectivity for each pair-wise connection was calculated as Pearson correlation between the two amplitude envelopes. Finally, the functional connectivity matrix was normalized using Fisher's r-to-z transformation.

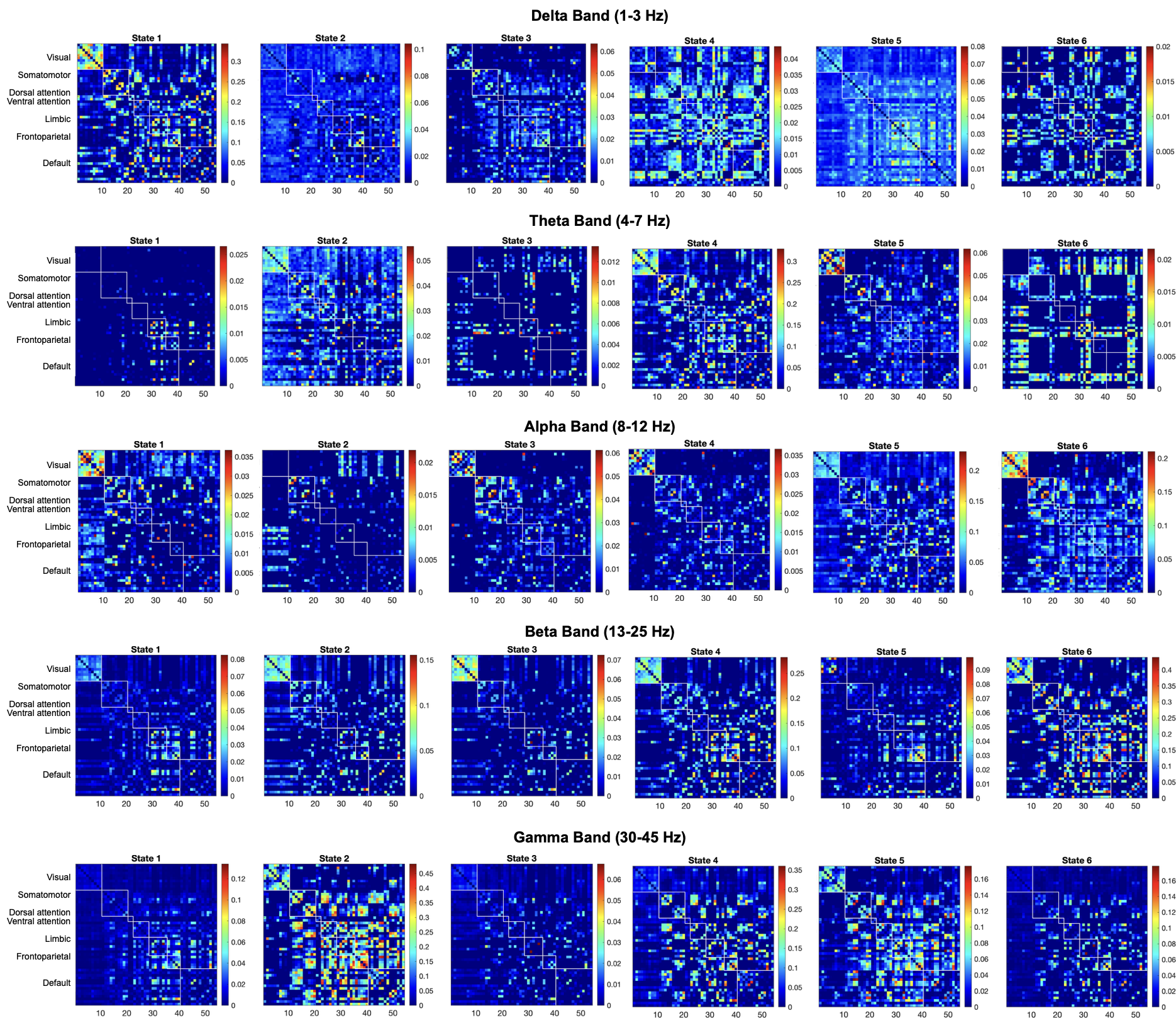

##### Figure S4. Distinct HMM connectome states for each canonical band.

HMM estimates the connectome states from the leakage-corrected EEG signals that are common to all subjects. The states are represented by z-scored functional connectivity (FC) matrices. The rows and columns represent 54 regions organized according to their membership to canonical intrinsic connectivity networks (ICNs) (Yeo et al., 2011).

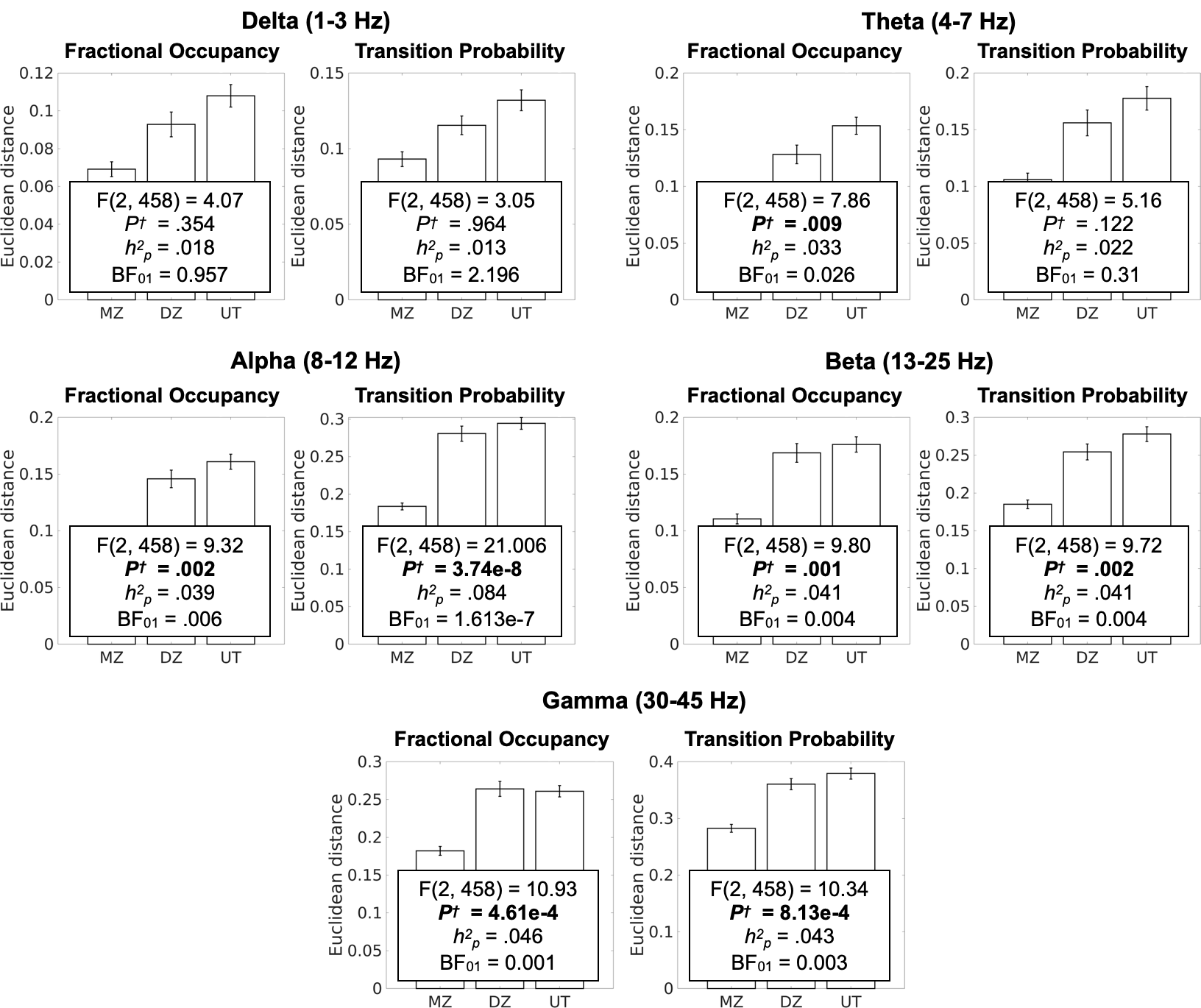

##### Figure S5. Heritability of temporal features of the dynamic connectome (*K* = 4).

Heritability of each of the multivariate features was assessed separately by one-way ANCOVAs of the factor sibling status (three levels: monozygotic twins (MZ), sex-matched dizygotic twins (DZ), and sex-matched pairs of unrelated individuals), adjusted for age and sex. The main effect of sibling status indicates the heritability, or genetic effect. (A) Bar graphs show that more genetically similar subject pairs have smaller Euclidean distance, which denotes similarity of a given temporal connectome feature. Specifically, temporal connectome features were more similar among MZ twin pairs than DZ twins, followed by pairs of unrelated individuals. P^†^: P values Bonferroni-corrected for 20 tests (four dependent variables and five frequency bands), h^2^_p_: Partial eta squared effect size.

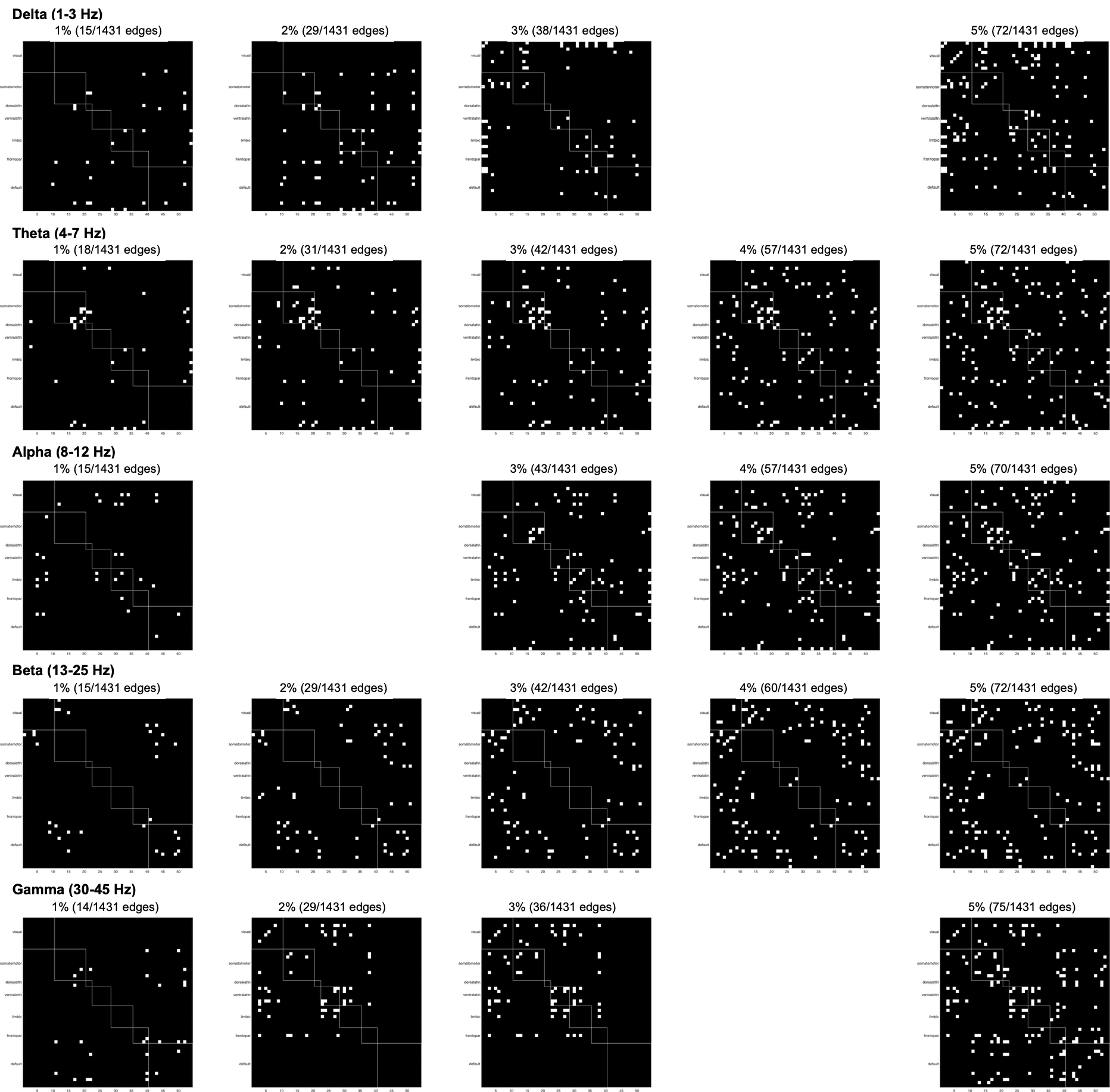

##### Figure S6. Binary NBS (data-driven) connectivity patterns.

We performed network-based statistics (NBS) to select connected sets of connections (i.e., clusters), in a data-driven manner for subsequent heritability analysis. We started with the absolute functional connectivity (FC) matrices, where all connections were transformed into absolute values to focus on the strength of connections, regardless of their connectivity direction. For each connection, an ANCOVA of the factor state was calculated to quantify the degree to which each connection’s connectivity strength differed across states, adjusted for age and sex. The ensuing connection-wise *F* statistics was threshold at arbitrary values, allowing 1 ~ 5% of connections to survive. NBS was applied to the threshold matrices, resulting in one significant cluster at each threshold visualized as symmetric binary matrices. We averaged the absolute FC values of all connections in the cluster for each state, together constructing the multivariate spatial feature.
